# A novel iPSC model of Bryant-Li-Bhoj neurodevelopmental/neurodegenerative syndrome demonstrates the role of histone H3.3 in chromatin dynamics, neuronal differentiation, and maturation

**DOI:** 10.1101/2024.08.26.609745

**Authors:** Annabel K. Sangree, Rajesh Angireddy, Janardhan P. Bhattarai, Yingqi Wang, Laura M. Bryant, Elisa A. Waxman, Dana E. Layo-Carris, Emily E. Durham, Kaitlin A Katsura, Emily E. Lubin, Xiao Min Wang, Kelly J. Clark, Minghong Ma, Elizabeth J. Bhoj

## Abstract

**Background:** Bryant-Li-Bhoj neurodevelopmental syndrome (BLBS) is neurogenetic disorder caused by variants in *H3-3A* and *H3-3B,* the two genes that encode histone H3.3. Ninety-nine percent of individuals with BLBS show developmental delay/intellectual disability, but the mechanism by which variants in H3.3 result in these phenotypes is not yet understood, limiting the therapeutic interventions available to individuals living with BLBS.

**Methods:** Here, we investigate how one BLBS-causative variant, *H3-3B* p.Leu48Arg (L48R), affects neurodevelopment using an induced pluripotent stem cell (iPSC) model differentiated to 2D neural progenitor cells (NPCs), 2D forebrain neurons (FBNs), and 3D dorsal forebrain organoids (DFBOs). We employ a multi-omic approach in the 2D models to quantify the resulting changes in gene expression and chromatin accessibility. We used immunofluorescence (IF) staining to define the identities of cells in the 3D DFBO and whole-cell patch clamp to investigate the electrophysiological properties of neurons in DFBOs.

**Results:** In the 2D systems, we found dysregulated gene expression and chromatin accessibility affecting neuronal fate, adhesion, neurotransmission, and excitatory/inhibitory balance. Immunofluorescence of DFBOs corroborated altered proportions of radial glia and mature neuronal populations. Patch clamp recordings revealed decreased electrical activity in neurons from L48R DFBOs compared to control DFBOs.

**Conclusions:** These data provide the first mechanistic insights into the pathogenesis of BLBS from a human-derived model of neurodevelopment, which suggest that H3.3 L48R increases *H3-3B* expression, resulting in the hyper-deposition of H3.3 into the nucleosome which underlies changes in gene expression and chromatin accessibility. Functionally, this causes dysregulation of cell adhesion, neurotransmission, and the balance between excitatory and inhibitory signaling. These results are a crucial step towards preclinical development and testing of targeted therapies for this and related disorders.

**Graphical Abstract:** 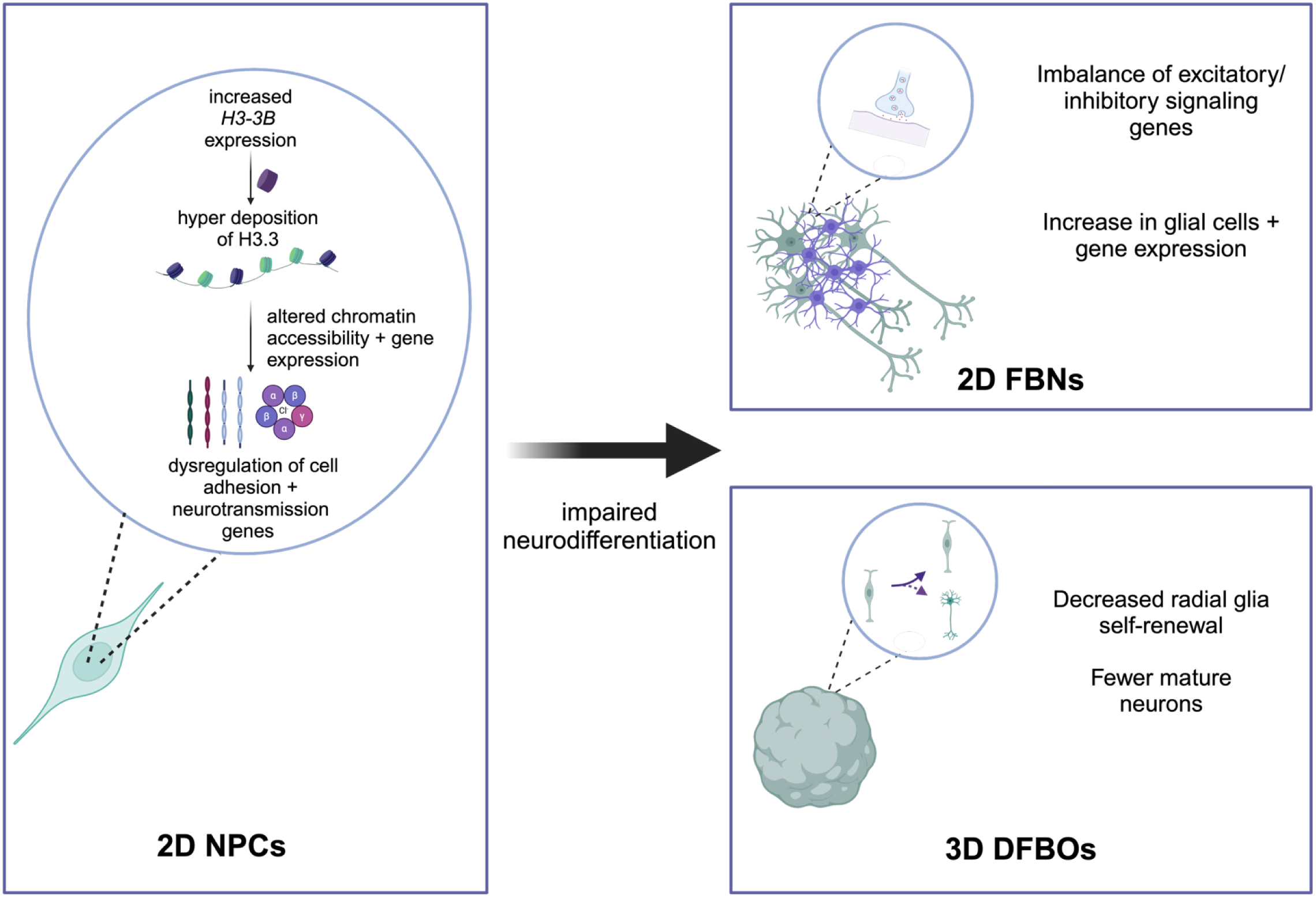

## BACKGROUND

Neurodevelopmental disorders (NDDs) affect 1-8% of the population and are often the result of aberrant components of epigenetic machinery following a genetic insult in the earliest stages of neurodevelopment^1^. Mendelian disorders of epigenetic machinery (MDEMs), caused by germline variants in 85 genes encoding histone writers, readers, erasers, and chaperones, are associated with a wide spectrum of NDDs, highlighting the brain’s vulnerability to transcriptional dysregulation^2^. Recently, variants in histones themselves emerged as causative in NDDs with the identification of *de novo* germline variants in the H4 encoding gene cluster^3,4^ and the H3.3-encoding genes *H3-3A* and *H3-3B*^5–7^.

Bryant-Li-Bhoj neurodevelopmental syndrome (BLBS, Online Mendelian Inheritance in Man (<u>OMIM</u>) #619720, #619721) is an NDD caused by heterozygous germline variants in *H3-3A* or *H3-3B*. Variants underlying BLBS occur throughout the coding sequences of *H3-3A* and *H3-3B* and are typically observed in just one individual^5,8^ ^(add^ ^in^ ^Okur^ ^citation)^. Phenotypically, affected individuals present almost universally with neurologic and craniofacial features. The most prevalent neurological feature is developmental delay/intellectual disability, which impacts 99% of published individuals with 51% of individuals exhibiting delayed or no attainment of sitting, 79% of individuals exhibiting delayed or no attainment of independent walking, and more than 40% of individuals not yet having attained their first word at >20 months^8^. From a neuroradiological standpoint, individuals most commonly have delayed myelination or hypomyelination, dysgenesis of the corpus callosum, and/or dilated ventricles^9^. As some individuals age, they develop an abnormal gait or progressive gait ataxia phenotype^5,7,8^. While no targeted therapeutics are available at this time, these neurodegenerative phenotypes suggest that BLBS could be amenable to therapeutic intervention to halt disease progression.

A mechanistic link has not yet been established between the physiologic function of H3.3 and its dysregulation in BLBS. H3.3 is a replication-independent histone that packages DNA and regulates gene expression. It is preferentially deposited by the chaperones DAXX or HIRA at enhancers, promoters, and gene bodies^10–18^. Because its deposition is not coupled to the cell cycle, H3.3 plays a critical epigenetic role in post-mitotic neuronal populations in the brain, especially in regulating transcription during learning and memory^19^. During early neurodevelopment in mice, H3.3 accounts for an estimated 31% of the total H3 pool^20^ and becomes the highly dominant form of H3 (>93%) in adult neurons^21^. Throughout all stages of neurodevelopment, H3.3 plays a crucial role in establishing chromatin landscapes, transcriptome, molecular identity, and axon connectivity in newly post-mitotic neurons^10^.

The role of H3.3 in disease was first appreciated in cancer through the discovery of somatic ‘oncohistone’ mutations^22–24^. While extensive work has been done to elucidate the mechanism of somatic oncohistone mutations like Gly34Arg/Val, the impact of germline missense variants, on neurodevelopment remains poorly understood..

Past studies into the effects of germline missense perturbation of H3.3 utilized cells from affected individuals, overexpression systems, and mouse models^5,6,29^. Dermal fibroblasts from affected individuals hyper-proliferate compared to age- and sex-matched controls^5^. *In silico* predictions suggest that certain mutants alter phosphorylation, glycosylation, methylation, acetylation, and ubiquitination histone post translational modifications (hPTMS) *in cis*^7^. In mice, three different variants at the same residue (p.Gly34Arg/Val/Trp) cause distinct developmental defects. p.Gly34Arg mice have severe neurodegeneration, p.Gly34Trp mice have obesity and bladder obstructions, and p.Gly34Val mice have intermediate phenotypes^29^. These models capture the same phenotypic heterogeneity recently reported amongst individuals living with BLBS, including those who share the same causative nucleotide level substitution^8^. These studies highlight the importance of studying individual variants, as both affected individuals and models display variable phenotypes. Moreover, there is a gap in knowledge as to how these variants affect human development in affected organs, most strikingly the brain, highlighting the necessity of interrogating the effect of BLBS-causing variants in human iPSCs.

Here, we profile the transcriptomic, chromatin accessibility, and cellular identity changes induced by one variant observed in our cohort, *H3-3B* p.Leu48Arg (L48R). This variant lies just beyond the N terminal tail, in the histone core^8^, a region of H3.3 that is less well studied in the context of disease. While the WT leucine has a hydrophobic side chain, arginine has a positively charged side chain. This L48R substitution is predicted to increase the distance between itself and other residues on H3.3, perhaps subtly opening the nucleosome at this position (Fig S1A). The individual harboring the L48R variant presents with neurologic features that include developmental delay/intellectual disability, microcephaly, hypomyelination, hypotonia that transitioned to hypertonia, and seizures.

To elucidate how this variant affects neurodevelopment, we utilized 2D and 3D models, allowing for both a detailed molecular analysis of gene expression and chromatin accessibility, as well as a platform to study the structural and cellular dynamics of BLBS in a more physiologically relevant context. In our 2D models of neural progenitor cells (NPCs) and forebrain neurons (FBNs) we elucidated global dysregulation in gene expression and chromatin accessibility of genes important for cell adhesion, axon development, and neurotransmission.

The 3D organoid models, which faithfully recapitulate developing neuroectoderm with the radial organization of NPC cells into structures that recapitulate the ventricular zone (VZ), subventricular zone (SVZ), and cortical plate (CP)^30^, recapitulated our findings of altered differentiation capacity of progenitor cell types Taken together, we find that H3.3 L48R perturbs neural differentiation and neuronal maturation in this iPSC model of brain development. Our results lay the foundation for future studies on BLBS and therapeutic interventions.

## METHODS

### Maintenance, editing, and differentiation of 2D cells

#### iPSCs

iPSCs were edited by the CHOP iPSC core as described^31^. The cells were single cell cloned and frozen at an early passage. Edited and isogenic control iPSCs were maintained in mTeSR™ Plus (STEMCELL Technologies 100-02 76) on 10cm plates coated with 1ug Matrigel® hESC-Qualified Matrix (Corning 354277) and three technical replicates were collected at passage 30 for RNA-sequencing.

#### NPCs

Cells were differentiated to NPCs using STEMdiff™ SMADi Neural Induction Kit following the monolayer culture protocol (STEMCELL Technologies 08582). Briefly, for each well, 2 million passage 31 iPSCs were resuspended in STEMdiff™ Neural Induction Medium + SMADi + 10 µM Y-27632 and plated in each well of a 6-well plate coated with 1ug of Matrigel. Cells well given a full medium change with STEMdiff™ Neural Induction Medium + SMADi except on days when they were passaged (approximately every seven days). Cells were collected at passage four post differentiation for RNA-sequencing. At the end of differentiation, cells were cryopreserved in STEMdiff™ Neural Progenitor Freezing Medium. Three separate differentiations were conducted by two different investigators to control for inter-differentiation variability.

#### FBNs

Two biological replicates of NPCs (passage 4) were thawed and differentiated to forebrain neurons using STEMdiff™ Forebrain Neuron Differentiation Kit (STEMCELL Technologies 08600). Briefly, NPCs were plated in PLO-Laminin coated 6 well plates (1.25 million cells per well) in STEMdiff™ Neural Induction Medium + SMADi, and after 24 hours, media was changed to STEMdiff™ Forebrain Neuron Differentiation medium. Cells were cultured in the same medium until they reached 80-90% confluency and passaged on to 12 well plates coated with PLO-laminin at a density of 175,000 cells per well in STEMdiff™ Forebrain Neuron Maturation medium. The FBNs were matured up to three weeks and collected for IF (2 biological replicates) and RNA-seq (3 technical replicates)

### Flow cytometry

Cells were harvested using Accutase, diluted with DPBS, and centrifuged at 3000 x*g*. They were then resuspended in 1.6% paraformaldehyde and incubated for 30 minutes at 37 °C with agitation. After a single wash with DPBS, the cells were resuspended in FACS Buffer (DPBS with 0.5% BSA and 0.05% sodium azide) and stored at 4 °C until flow cytometry analysis. For the flow cytometry analysis, cell permeabilization was performed with saponin buffer (diluted to 1× in water; Biolegend, 421,022), followed by a 1-hour incubation with antibodies diluted in saponin buffer: Forse-1 (DSHB, 1:100), NCAD (Cell Signaling #14215, 1:300), and an IgM isotype control (Santa Cruz, sc-3881, 1:100). The cells were then washed with either saponin buffer or FACS buffer and incubated for 1 hour with secondary antibodies: goat anti-rabbit Alexa 488 (Jackson ImmunoResearch, 111–545-144, 1:500) or goat anti-mouse IgM-conjugated Alexa 488 (Thermo Fisher Scientific, A21042, 1:500). Finally, the cells were washed again and resuspended in FACS buffer. Analysis was conducted using a CytoFLEX flow cytometer (Beckman Coulter) with the FlowJo software program (BD).

### RNA sequencing

#### RNA extraction, library prep & sequencing

Total RNA (3 replicates of each cell type) was extracted from iPSCs and NPCs using the Zymo Direct-zol RNA Miniprep (Cat # R2050 or from FBNs with the Zymo Direct-zol RNA Microprep (Cat # R2062). QC was performed using the Gx Bioanalyzer/Agilent Tapestation for traces and Qubit for concentration. Libraries were prepared using the Illumina Stranded Total RNA Prep, Ligation with Ribo-Zero Plus. 200ng of cDNA libraries were amplified using 12 cycles of PCR and sequenced on a v1.5_SP (200 cycle) flow cell with the NovaSeq6000. Reads were demultiplexed using Dragen Demultiplexing.

#### Data analysis

At least 15E6 million r eads were aligned to Homo_sapiens.GRCh38.cdna.all using kallisto^32^(v0.50.1). Replicates were averaged and differential expression was calculated using DeSeq2^33^(v1.44.0) in R v 4.4.0 using RStudio 2024.04.1+748. Volcano plots were made using EnhancedVolcano^34^(v1.22.0). We defined genes using adjusted p-values (Benjamini and Hochberg corrected) and log2fold changes output by DESeq2 with significantly altered expression with an adjusted p-value ≤ 0.05 and log fold change with an absolute value of ≥ 1. GO analyses were performed using clusterProfiler^35^ (v4.12.0) using biological pathway terms. Revigo^36^ was used to summarize GO terms and these terms were clustered using a distance matrix and hierarchical clustering. Each cluster was labeled with one representative term, obtained by selecting the term within each cluster with the largest LogSize, the Log10 value of the number of annotations for GO Term ID in selected species in the European Bioinformatics Institute Gene Ontology Annotation database. The list of astrocyte specific genes used to analyze the FBN data was obtained from PanglaoDB^37^, filtered for human, and a specificity score ≤ 0.5 (meaning that these genes are expressed in cells other than astrocytes less than or equal to 50% of the time).

### Western Blots

Histones were extracted using the Abcam Histone Extraction Kit (ab113476). Protein concentration was quantified using the Pierce™ BCA Protein Assay Kit (A65453) and run using the SkanIt software v6.1 by measuring the absorbance at 550nm. 2ug of histone extract was loaded per reaction and separated by electrophoresis using NuPAGE^TM^ 4-12% Bis-Tris Protein Gel (ThermoFisher Scientific). Proteins were transferred to a polyvinylidene difluoride (PVDF) membrane using an XCell SureLock Mini-Cell Electrophoresis System (ThermoFisher Scientific). The membrane was blocked using 5% dry milk in Tris-Buffered Saline (Bio Basic, Markham) with Tween-20 for 1 hour at room temperature. After briefly rinsing with Tris-Buffered Saline (Bio Basic, Markham) with Tween-20, the membrane was incubated overnight at 4°C with a previously validated primary antibody (Table 5). The membrane was then washed with Tris-Buffered Saline with Tween-20 and incubated in secondary antibody diluted 1:10,000 (Goat Anti-Rabbit IgG Polyclonal from LI-COR Biosciences) for 1 hour at room temperature. Blots were visualized using the ODYSSEY® DLx (LI-COR Biosciences) and quantified using ImageJ.

### ATAC-sequencing

#### Library preparation & sequencing

ATAC-seq libraries were generated using the ATAC-seq Kit from Diagenode (Diagenode, A Hologic Company) according to manufacturer instructions. Briefly, cells were centrifuged at 500 × g for 10 minutes using a fixed angle refrigerated centrifuge, followed by a wash with 500 μl of cold 1x PBS and a second centrifugation at 500 × g for 10 minutes. Cells were counted and approximately 50,000 cells were lysed using cold ATAC lysis + Digitonin (2%). Immediately after lysis, cell nuclei were spun again at 500 × g for 10 minutes and separated from the lysis buffer. Immediately following the nuclei extraction, the pellet was resuspended in transposase reaction mix. The transposition reaction was carried out for 30 minutes at 37 °C. Directly following transposition the samples were purified using the kit’s provided columns. Following purification, library fragments were amplified using 1x Amplification master mix and the following PCR conditions: 72°C for 5 minutes, 98°C for 30 seconds, followed by thermocycling at 98°C for 10 seconds, 63°C for 30 seconds and 72°C for 1 minute for 5 cycles, in this step Unique Dual Indexes Primer Pair were incorporated for multiplexed sequencing. To reduce amplification bias after the first 5 cycles, qPCR was used to determine the number of PCR cycles needed to produce the concentration required for sequencing. Each pre-amplified library was further amplified for 6 cycles as determined by the qPCR step. Final libraries were bead purified (Beckman Coulter), then assessed for size distribution and concentration using the Bioanalyzer High Sensitivity DNA Kit (Agilent Technologies). The resulting libraries were pooled in equal molarity, the pool was diluted to approximately 2 nM using 10 mM Tris-HCl, pH 8.5, denatured, and loaded onto a P1-100 (2×50) flow cell on an Illumina NextSeq 6000 (Illumina, Inc.) according to the manufacturer’s instructions. De-multiplexed and adapter-trimmed sequencing reads were generated using bcl2fastq.

#### Data processing and analysis

Reads were successfully aligned to *Homo* sapiens genome build Genome Reference Consortium Human Build 38 (GRCh38) with botwie2^38^ (v2.5.4). PCR duplicates were removed using picard^39^ (v3.2.0), mitochondrial regions were removed, and blacklisted regions were identified using ENCODE (GRCh38_unified_blacklist.bed) and removed. Replicates were merged and subset to 55E6 reads for both Control and L48R NPCs. Peaks were called using macs3^40^ (v3.0.1) with a q-value cutoff of 0.05. Peaks were annotated using ChIPSeeker^36^(v1.40.0). DiffBind^69^(v3.14.0) was used to determine differential accessibility between L48R and control peaks and plotted using the DiffBind profile plot. For transcription factor footprinting, reads were shifted to account for the Tn5 overhang using ATACshift from deeptools^44^, and motifs were identified using ATACseqTFEA^55^ with human motifs “best_curated_Human.rds”. This packages collects motifs from jasper2018, jolma2013 and cisbp_1.02 from package motifDB (v 1.28.0) and merged by distance smaller than 1e-9 calculated by MotIV::motifDistances function (v 1.42.0). Genome browser plots were made using IGV(v2.17.4)

### qPCR validation

Total RNA was extracted from iPSC and NPCs using the Zymo Direct-zol RNA Miniprep (Cat # R2050 or from FBNs with the Zymo Direct-zol RNA Microprep (Cat # R2062). The concentration was quantified using the NanoDrop 8000 UV-Vis Spectrophotometer (Thermo Scientific). cDNA was synthesized from 1ug of RNA using Superscript IV (Thermo Scientific). RT-qPCR reactions were set up using cDNA, 2X PowerUp™ SYBR® Green Master Mix (Applied Biosystems), and gene specific primers using GAPDH as an internal control. Samples were run in a 96-well plate on the QuantStudio™ 3 System using the Comparative Cт (ΔΔCт) method. The primers used are listed in Table 6. Two representative primer pairs are presented below:

*GAPDH* F: GGCATCCTGGGCTACACTGA, R: GAGTGGGTGTCGCTGTTGAA
*H3-3B* F: TCGTTGGGCGGTGCTGGTTTTT R: TTAGGCCACTTCTCCGACCGCC

### Dorsal Forebrain Organoids (DFBOs)

Forebrain organoids were generated using STEMdiff™ Dorsal Forebrain Organoid Differentiation Kit (Cat: 08620) according to manufacturer instructions, and this protocol was adopted from^45^. Briefly, embryoid bodies (EB) were generated in AggreWell™800 24-well plate in a seeding medium with 10,000 cells per microwell. This method will generate 500 equalized EBs per one well of a 24-well plate. After 24 hours, the media was changed to a Forebrain Organoid formation medium, and the EB was maintained for five days. EBs were transferred to ultra-low attachment 6-well plates (40 organoids per well) in organoid expansion medium. Organoids were maintained in the same medium for a period of 25 days with frequent media changes. After the neuroectoderm expansion, the media was changed to the Forebrain Organoid Differentiation medium and maintained organoids until day 43 with frequent media changes. For long-term maintenance of DFBO, the media was changed to a Forebrain Organoid maintenance medium. For the analysis, we harvested organoids at day 25 (Neuroectoderm expansion) and day 62 (DFBO maturation) and day 90 (Electrophysiology).

#### Embedding

Organoids were washed with D-PBS then fixed in 5mL fresh 4% paraformaldehyde (PFA) solution and incubated overnight at 4°C. Organoids were washed in PBS-T and stored in PBS-T at 4°C for 1 week. Organoids were then incubated in 30% sucrose solution and equilibrated overnight at 4°C. Equilibration was complete once organoids were no longer floating in the 30% sucrose solution. Organoids were then incubated at 37°C in porcine gelatin for 1 hour and embedded in this solution in embedding molds. Six organoids were embedded per block and allowed to solidify at RT and 4°C before being snap frozen in a dry ice/ethanol slurry (dry ice + 100% ethanol). Frozen samples were stored at −80°C until ready for cryosectioning.

#### Cryosectioning and Immunoflourescence Staining

Blocks were removed from the −80°C freezer and allowed to equilibrate to −26°C in the cryostat (Microm HM 505E, CHOP Pathology Core) for at least 30 minutes before sectioning. Sections were cut at 16um using a high-profile blade (Leica Biosystems 14035838926) and mounted on colorfrost Plus Microscope Slides (Fisher Scientific 12-550-17). Slides were stored at −80C until staining. Slides were allowed to come to room temperature before staining. Hydrophobic barriers were drawn around sections. Antigen retrieval using sodium citrate pH 6.0, was performed for the CTIP2 staining. Slides were steamed in a food steamer for 20 minutes in citrate buffer, washed with PBS-T at 37°C for 10 minutes to fully remove gelatin from slides, blocked in a humidified chamber for 1 hour, and incubated with primary antibody overnight. The next day, primary antibody solutions were removed, slides were washed 3 times with PBS-T and incubated in secondary antibody at room temperature in a humified chamber for 2 hours. After incubation, secondary antibody solution was removed, slides were washed 3 times with PBS-T for 30 minutes, allowed to air dry for 5 minutes, then one drop of PermaFluor™ was added to slides to add coverslip.

#### Image acquisition and quantification

Images for quantification were acquired using a Keyence BZ-X800 (Itasca, IL) or the Leica STELLARIS 5 Confocal Microscope. 10x images taken of DFBOs were stitched together using Keyence BZ-X Series Analysis Software. FIJI (v 2.14.0/1.54f) was used to quantify the fraction of cells that were positive for cell fate markers. Images were separated by channel, converted to binary, then particles were analyzed for each image. Positive cells for each marker were normalized to the total number of cells counted (DAPI) in each image.

### Patch Clamp Electrophysiology

Dorsal forebrain organoids at maturation day 90 were prepared for electrophysiological recordings. Before recording, DFBOs were transferred to a recording chamber and held by a nylon slice holder and continuously perfused with oxygenated artificial cerebrospinal fluid (ACSF) containing the following (in mM): 126 mM NaCl, 2.5 mM KCl, 1.2LmM MgSO_4_, 2.4LmM CaCl_2_, 25LmM NaHCO_3_, 1.4LmM NaH_2_PO_4_, 11 mM glucose, and 0.6 mM sodium L-ascorbate) and continuously bubbled with 95% O_2_ and 5% CO_2_). Neurons were visualized with a 40× water-immersion objective under an Olympus BX51WI upright microscope equipped with epifluorescence. Recording pipettes were made from borosilicate glass (GC210F-10; Harvard Apparatus) with a Flaming-Brown P-97 puller (Sutter Instruments; tip resistance 5–8 MΩ). The pipette solution contained the following (in mM): 120 K-gluconate, 10 NaCl, 1 CaCl2, 10 EGTA, 10 HEPES, 5 Mg-ATP, 0.5 Na-GTP, and 10 phosphocreatine. Whole-cell patch-clamp recordings were conducted in both current and voltage-clamp modes. Electrophysiological recordings were controlled by an EPC-10 amplifier combined with Pulse Software (HEKA Electronic) and analyzed using Igor Pro.

For a subset of neurons, intrinsic membrane properties were measured via current injections under current-clamp mode. The resting membrane potential was determined when there was no current injected. For the remaining properties, the cell membrane potential was kept at −70 mV. The input resistance was calculated as the slope of the current-voltage (I-V) curve (current steps from −100 pA with 10 pA increments). The action potential (AP) threshold was determined as the point with the maximum curvature before the AP evoked by the minimum current step.

### Statistics

Comparisons were made between Control and L48R cells for each timepoint. For RT-qPCR and IF, statistics were calculated using Welch Two Sample t-test in R v 4.4.0 using RStudio 2024.04.1+748. A p-value ≤ 0.05 is denoted with *, p-value ≤ 0.01 is denoted with **, and p-value ≤ 0.001 is denoted with ***. For RNA-seq, statistics were calculated using DESeq2^33^, and for ATAC-seq statistics were calculated using DiffBind^43^. No statistical method was used to predetermine sample size.

## RESULTS

### L48R variant affects iPSC differentiation into 2D neural progenitor cells (NPCs) and 2D forebrain neurons (FBNs)

We introduced the heterozygous c.146T>G *H3-3B* Leu48Arg (L48R) variant into CHOPWT14 iPSCs (from a Hispanic 46XX donor) using CRISPR/Cas9 homology directed repair (HDR)^31^ followed by single cell clone selection and validation by Sanger sequencing (Fig. 1A, Fig. S1A-C). To validate pluripotency, we performed RT-qPCR for *OCT4* and *NANOG* in these iPSCs (Fig. 1B). We did not observe significant changes in the expression of these markers between L48R and control cells, confirming that this variant did not have a large impact on the identity of iPSCs.

**Figure 1.**
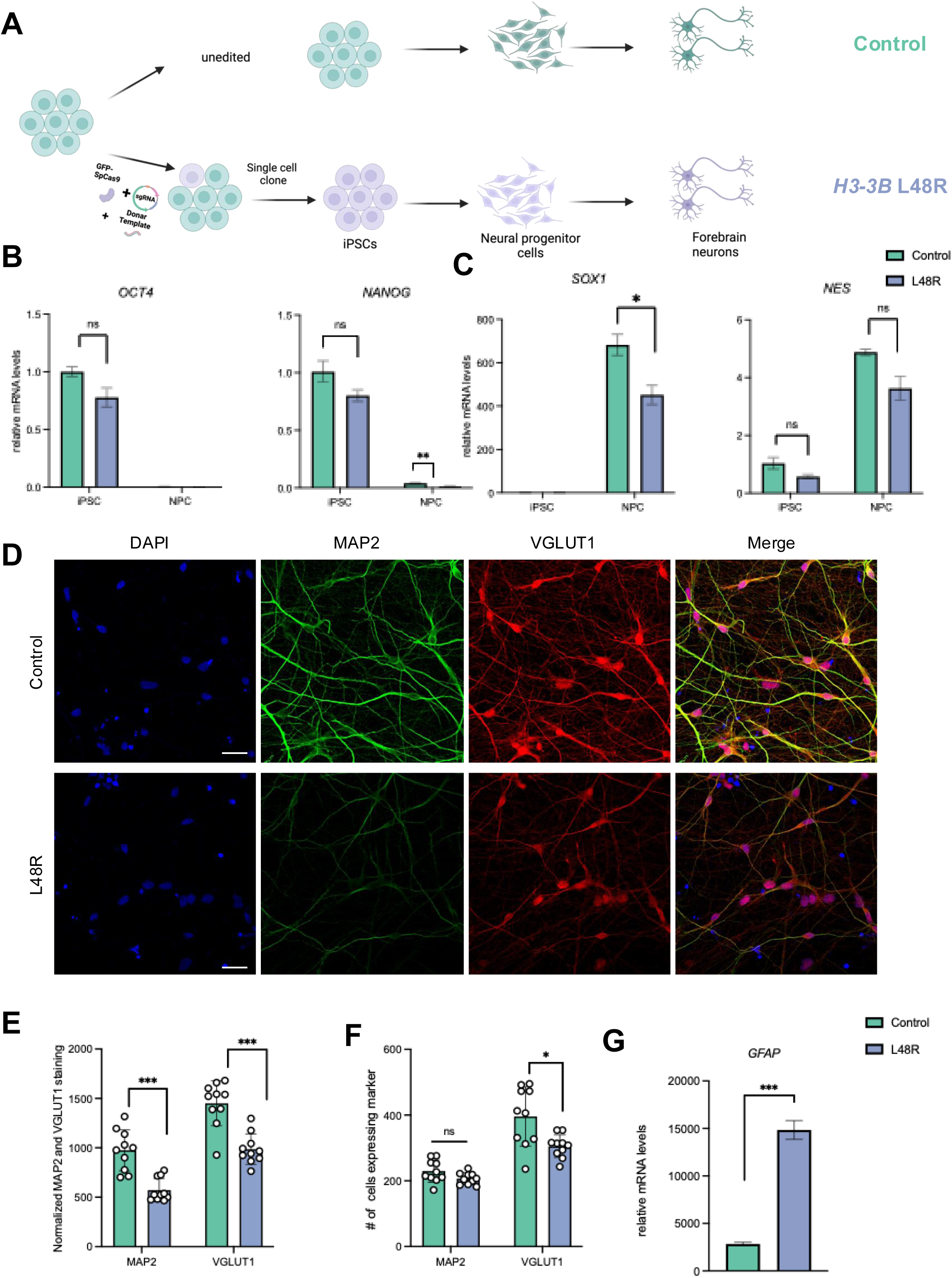
Generation and initial characterization of L48R 2D iPSCs, NPCs and FBNs. A) c.146T>G *H3-3B* Leu48Arg (L48R) variant was introduced to CHOPWT14 hiPSCs using CRISPR HDR and pure population of edited cells were attained and differentiated into NPCs and FBNs. B-G) **L48R and isogenic control iPSCs showed no significant differences in pluripotency markers (OCT4, NANOG). L48R NPCs had reduced expression of NANOG (**p = 0.00836) **and SOX2 (p = 0.02608) and FBNs exhibited impaired neuronal extension (MAP2, VGLUT1) and increased astrocytic marker GFAP (p = 1.104e-06)** D) Immunofluorescence of FBNs stained with DAPI, MAP2 (neuronal extensions), VGLUT1 (glutamatergic neurons). E-F) Quantification of images in D) using FIJI. n = 10 images from 3 replicates for each condition. MAP2 p = 9.536e-05, VGLUT1 p = 6.694e-05. G) RT-qPCR for astrocyte marker *GFAP* p = 1.104e-06. Comparisons made via t-test and shown with brackets. *p ≤0.05, ** p ≤0.01, *** p ≤0.001

To begin delineating the effects of the H3.3 L48R variant on brain development and neuronal function, we differentiated L48R and isogenic control iPSCs into NPCs using a dual SMAD inhibition protocol^46^. We found that the L48R NPCs had lower expression of NPC markers *NES* (encodes Nestin) and *SOX1* than control (Fig. 1C), indicating that this variant affects early neuronal differentiation. FACS analysis on control and L48R NPC stained with progenitor markers Forse1 and NCAD indicates that the L48R mutation has no effect on the overall NPC population (Fig. S1D-G). Next, we differentiated the NPCs to FBNs using a protocol that generates a mixed population of 90% forebrain-type neurons (MAP2+ and TUJ1+) with about 10% glial cells (GFAP+) (Stem Cell Technologies #08600, #08605). We immunostained FBNs at day 20 of maturation for the neuronal marker TUJ1 and glial marker GFAP. We observed decreased TUJ1 and increased GFAP intensity, however neither of these reached significance (Fig. S1H, I). Interestingly, there was no significant change in the total number of cells expressing TUJ1, and a decrease in L48R cells expressing GFAP compared to control (Fig. S1J). This suggests that the increased GFAP intensity might be due to an increase in reactive astrocytes, rather than total cell number. . Further staining indicated decreased intensity of MAP2+ neuronal extensions and glutamate transporter VGLUT1 in the L48R FBNs compared to isogenic controls (Fig. 1D, E). We found no change in the number of cells expressing MAP2, or relative *MAP2* mRNA expression measured via RT-qPCR between the two conditions (Fig. 1F, Fig. S1K). We found a significant decrease in VGLUT1 staining and a significant increase in *GFAP* mRNA levels in L48R FBNs compared to isogenic controls (Fig. 1F,G). Taken together, we observed disruptions in the differentiation of L48R iPSCs to NPCs, which impacted subsequent differentiation into FBNs.

### Transcriptomic changes in L48R Neural progenitor cells affect differentiation into 2D Forebrain Neurons

To understand the global transcriptomic changes induced by L48R at various stages of differentiation we performed bulk RNA-seq in 3 biological replicates of iPSCs, NPCs, and FBNs (Supplementary File 1). Principal component analysis (PCA) on the different cell types identified three distinct clusters based on cell identity (Fig. S2A). We identified 96 genes with significantly altered expression (p ≤ 0.05) in L48R iPSCs compared to the isogenic controls (Fig. 2A). Using an additional threshold of absolute value log2-fold change (LFC) ≥ 1, 37 genes were significantly upregulated, while 20 were downregulated. Using a gene ontology (<u>GO</u>) analysis, we found no significantly altered pathways in the L48R iPSCs compared to isogenic controls, but did identify the upregulation of *CDKL5* (LFC = 1.46, adjusted p-value = 6.36E-09) and *SCML2 (*LFC = 1.04, adjusted p-value = 4.98-E07) both of which are genes important for cell cycle progression and differentiation^47,48^ (Table 1). We validated this finding using RT-qPCR in iPSCs, NPCs and FBNS, and found the specific upregulation of *CDKL5* and *SCML2* in L48R cells at the iPSC stage only (Fig S2 B). These transcriptomic results indicate that L48R might impact the ability of iPSCs to proliferate and differentiate into NPCs, but it does not affect the identity of iPSCs.

**Figure 2.**
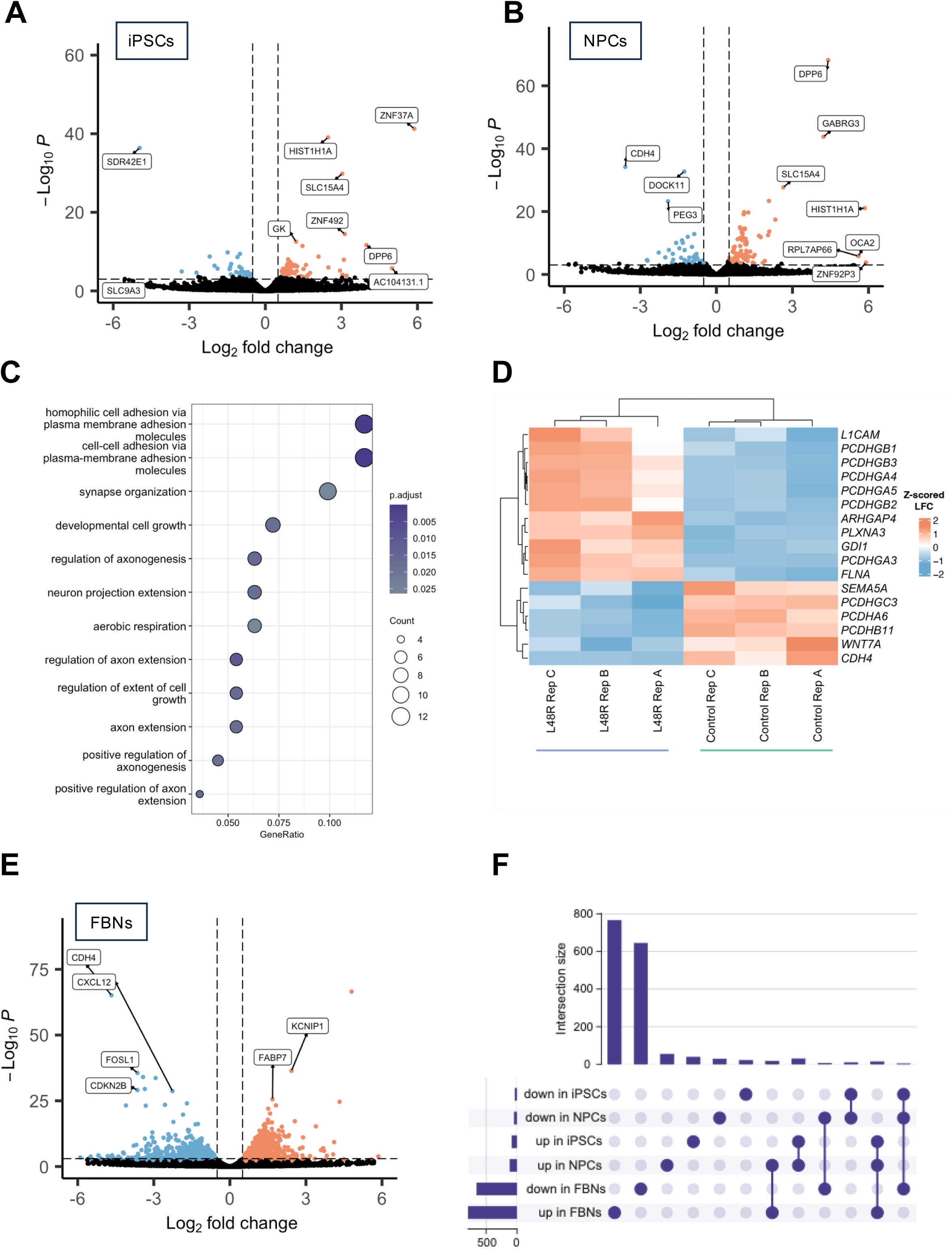
Transcriptomic changes in L48R progenitor cells affect differentiation into FBNs. A-C) **RNA-seq reveals dysregulation of cell adhesion (e.g., CDH4, WNT7A) and axonogenesis (e.g., PLXNA3) pathways in L48R NPCs.** Blue dots represent downregulated genes, red = upregulated genes. Dotted horizontal line denotes an adjusted p-value of ≤ 0.01 and dotted vertical lines indicate a log2fold change (LFC) with an absolute value of ≥ 1. **GO analysis identifies upregulation of genes affecting synapse activity and neurotransmitter uptake in FBNs, suggesting L48R affects both early differentiation and neuronal function.** D) Heatmap representing the Z-scored LFCs of the genes comprising cell adhesion and axonogenesis GO terms. These genes appear in 9/12 significant GO terms between L48R and control NPCs. Descriptions of function and disease association are in Table 1 for these genes. E) Volcano plot of L48R vs control FBNs formatted as in A-B. F) Upset plot of shared hits between the RNA-seq datasets generated in iPSCs, NPCs and FBNs. n = 2 commonly downregulated genes, n=10 commonly upregulated genes.

**Table 1.**
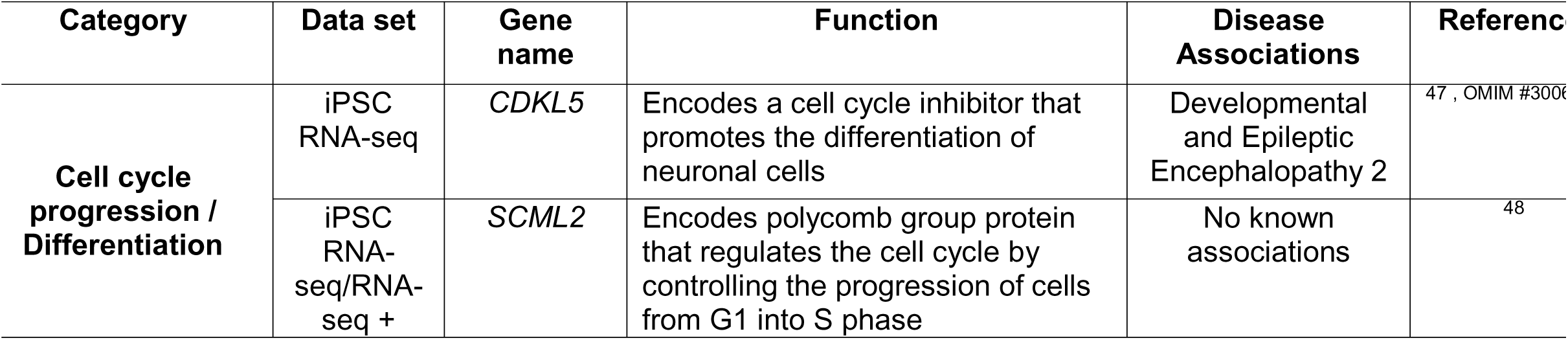
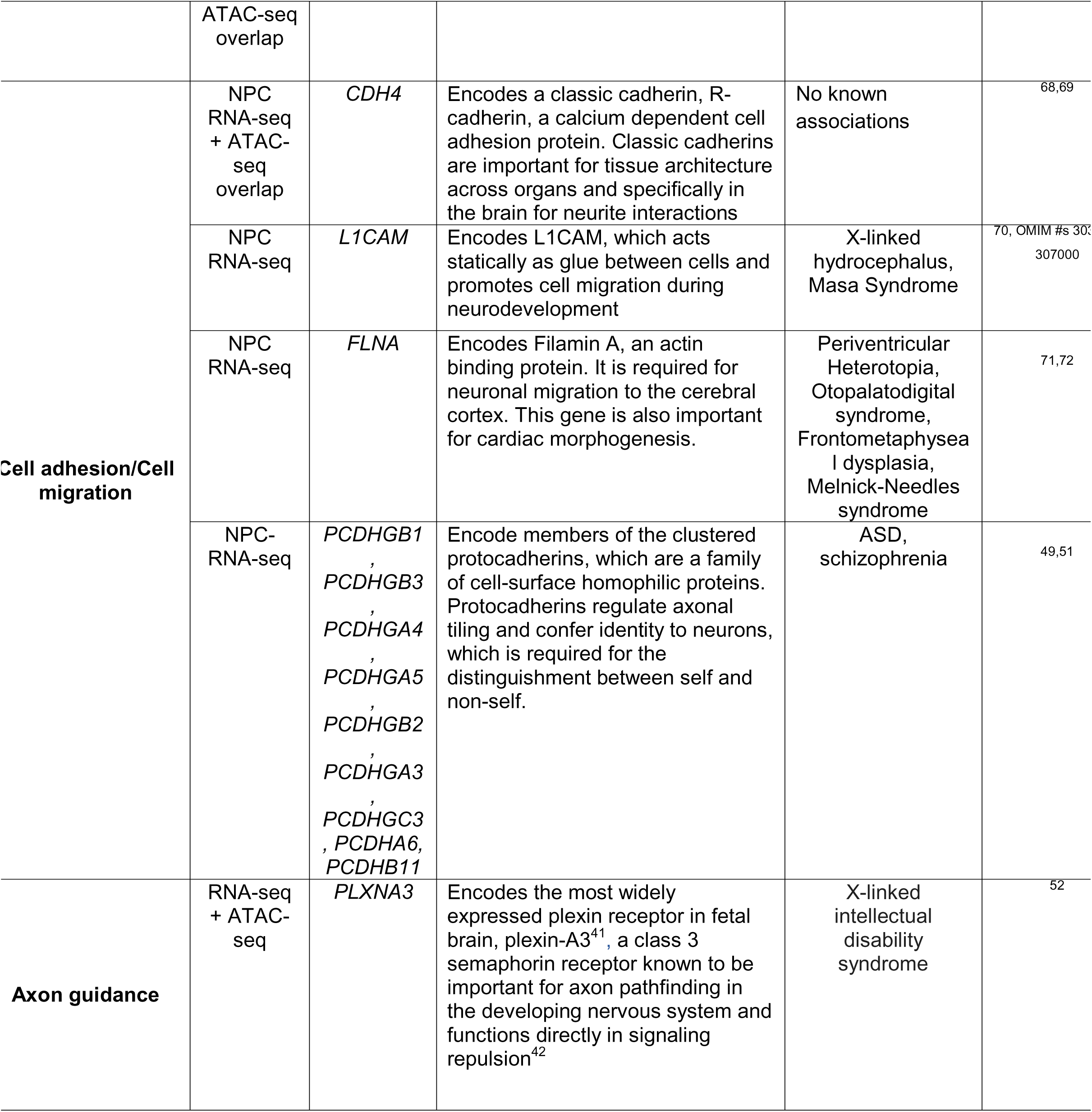
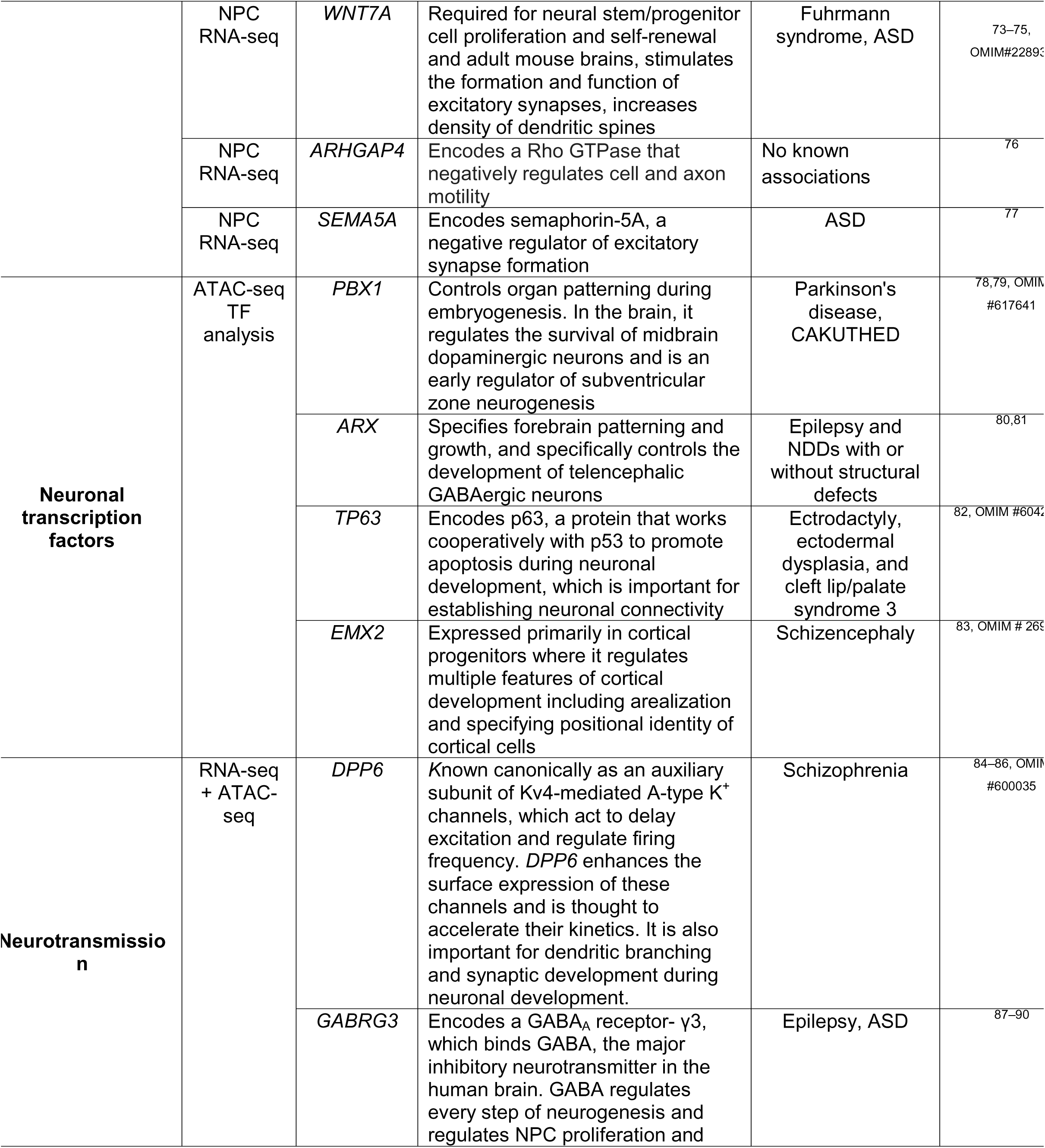
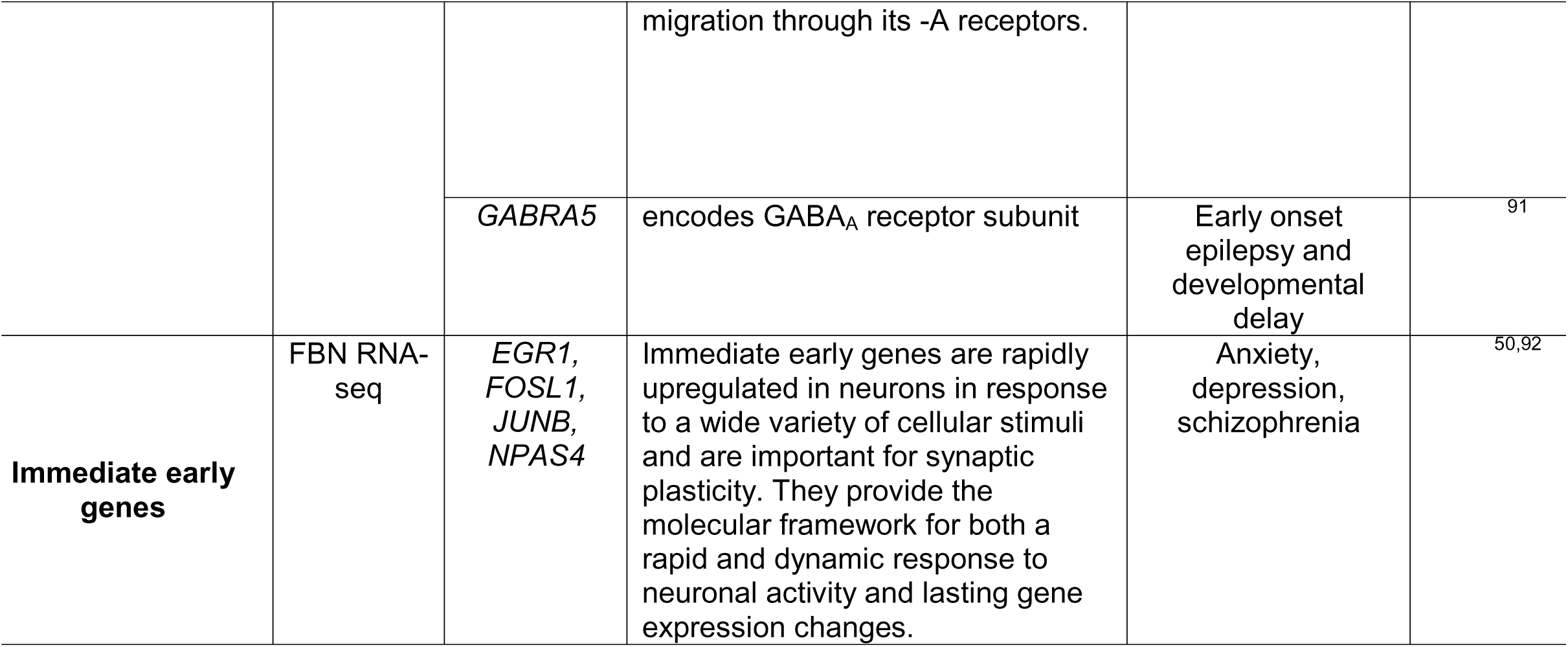
Function and disease associations of the dysregulated genes in L48R compared to isogenic controls.

Nearly twice as many genes were dysregulated in L48R NPCs. Bulk RNA-seq on NPCs identified 139 differentially expressed genes in L48R NPCs (p ≤ 0.05) compared to control NPCs. 53 genes were upregulated (LFC ≥ 1) and 27 downregulated (LFC ≤ −1) (Fig. 2B). GO analysis for all differentially expressed genes indicate that processes like cell migration and establishment of synapses are affected, including pathways involved in cell adhesion, synapse organization and regulation of axonogenesis, (Fig. 2C, Table 1). When focusing on genes comprising the cell adhesion and axonogenesis pathways most genes related to these processes were upregulated in L48R NPCs compared to controls (Fig. 2D)., Several protocadherin-encoding genes which mediate neurite repulsion^49^, were upregulated, while *CDH4* and *WNT7A* were downregulated, indicating an imbalance in cell adhesion and axon development cues (Table 1).

When L48R NPCs were converted to FBNs, we observed massive transcript-level dysregulation (773 genes up, 965 down) (Fig. 2E). *EGR1, FOSL1, JUNB,* and *NPAS4,* immediate early genes which mediate genetic responses to environmental cues^50^, were downregulated (Fig. 2E, Table 1). GO analysis identified the upregulation of terms related to synapse organization and activity, neurotransmitter uptake, and nervous system development (Fig. S2C). It also identified terms related to G/M cell cycle phase transition and mitotic cell cycle phase transition, which are notable given that neurons are post-mitotic (Fig. S2C). We also found that astrocyte-associated genes like *AQP4*, *SLC1A3,* and *SLC6A11* are upregulated in L48R FBNs, suggesting that the mixed population of cells were transciptomically more similar to proliferating astrocytes, as opposed to post-mitotic neurons (Fig. S2D). To this end, we also observed increased BrdU staining in L48R FBNs, indicative of proliferation in the L48R cells compared to controls following FBN differentiation (Fig. S2E). When we analyzed the GO terms for down-regulated genes in FBNs, we observed terms like regulation of MAPK cascade, regulation of apoptotic signaling pathway, and cellular response to external stimulus (Fig. S2F). Given the dysregulation of fundamental pathways that we observed in NPCs and the downregulation of immediate early genes, it is likely that the widespread dysregulation observed in the FBNs is secondary to these primary deficits.

To identify causative genes that might affect the overall differentiation of L48R cells, we looked for commonly dysregulated genes at the iPSC, NPC, and FBN stage (Fig. 2F). Two genes were commonly downregulated, and 13 were upregulated in all cell types (Table 2). Restricting the analysis to genes commonly dysregulated later in the differentiation proccess at the neuronal NPC and FBN stages, we identified 29 genes that were upregulated in both NPCs and FBNS, and 8 that were downregulated (Table 3). All of the genes dysregulated early in differentiation persisted throughout (Fig. S2G).

**Table 2.**
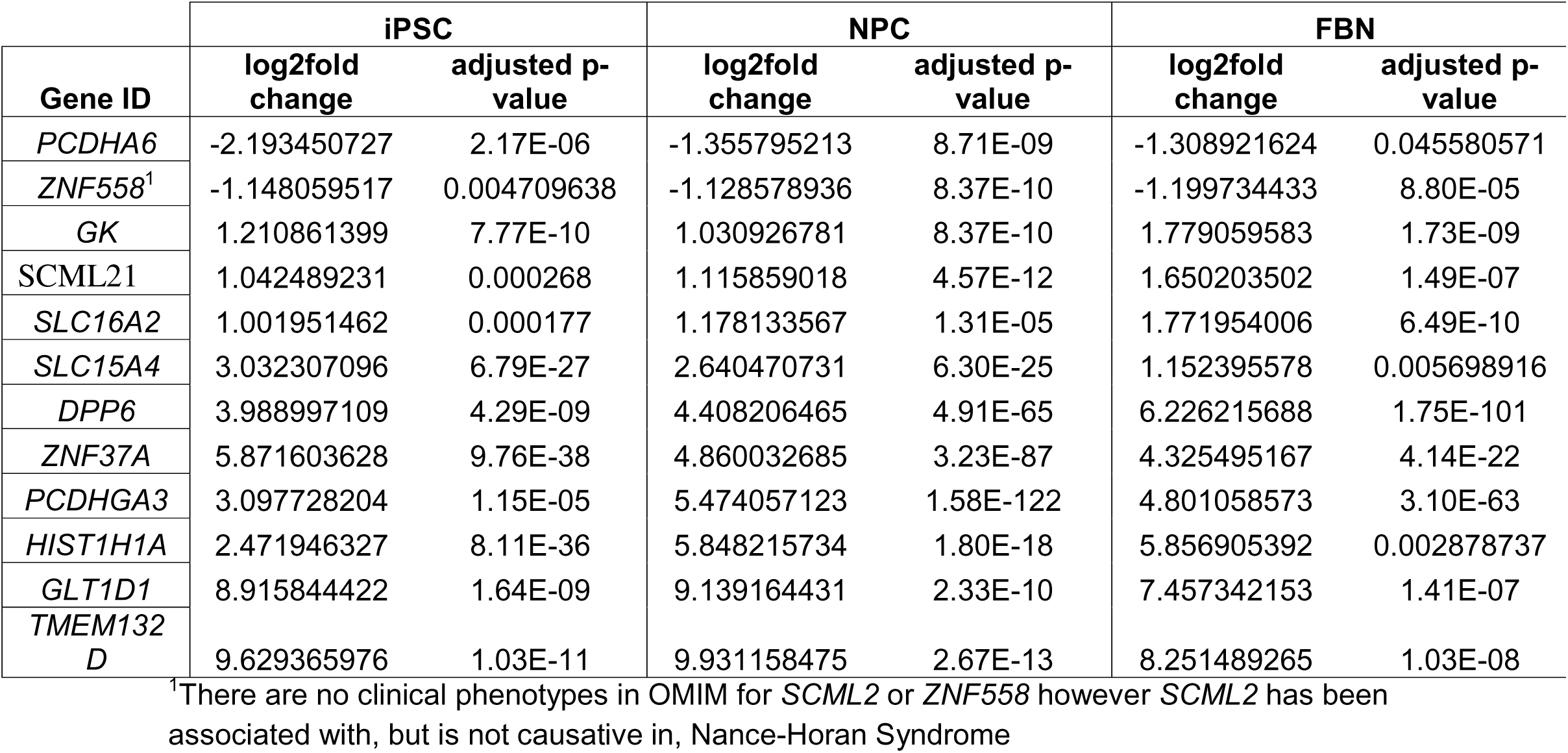
Genes dysregulated across 2D iPSCs, NPCs and FBNs.

**Table 3.**
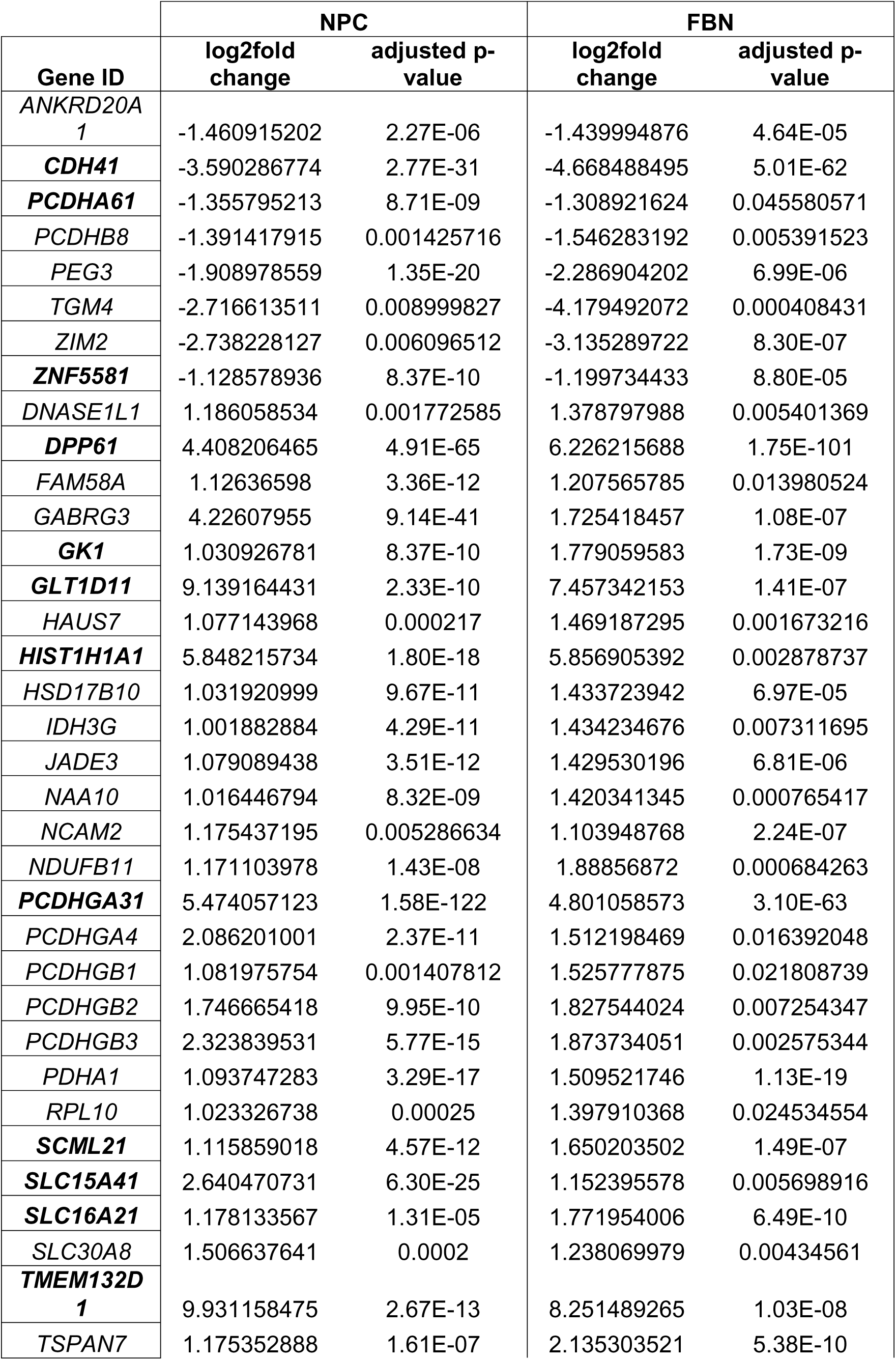
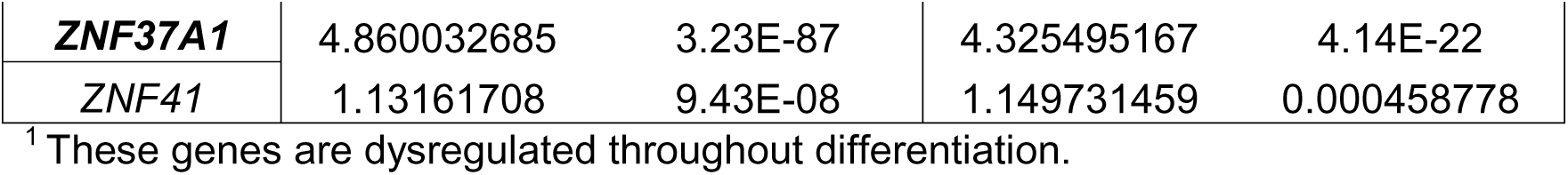
Genes dysregulated in both L48R NPCs and FBNs compared to isogenic controls.

|  | NPC |  | FBN |  |
| --- | --- | --- | --- | --- |
| Gene ID | log2fold change | adjusted p-value | log2fold change | adjusted p-value |
| *ANKRD20A1* | -1.460915202 | 2.27E-06 | -1.439994876 | 4.64E-05 |
| ***CDH41*** | -3.590286774 | 2.77E-31 | -4.668488495 | 5.01E-62 |
| ***PCDHA61*** | -1.355795213 | 8.71E-09 | -1.308921624 | 0.045580571 |
| *PCDHB8* | -1.391417915 | 0.001425716 | -1.546283192 | 0.005391523 |
| *PEG3* | -1.908978559 | 1.35E-20 | -2.286904202 | 6.99E-06 |
| *TGM4* | -2.716613511 | 0.008999827 | -4.179492072 | 0.000408431 |
| *ZIM2* | -2.738228127 | 0.006096512 | -3.135289722 | 8.30E-07 |
| ***ZNF5581*** | -1.128578936 | 8.37E-10 | -1.199734433 | 8.80E-05 |
| *DNASE1L1* | 1.186058534 | 0.001772585 | 1.378797988 | 0.005401369 |
| ***DPP61*** | 4.408206465 | 4.91E-65 | 6.226215688 | 1.75E-101 |
| *FAM58A* | 1.12636598 | 3.36E-12 | 1.207565785 | 0.013980524 |
| *GABRG3* | 4.22607955 | 9.14E-41 | 1.725418457 | 1.08E-07 |
| ***GK1*** | 1.030926781 | 8.37E-10 | 1.779059583 | 1.73E-09 |
| ***GLT1D11*** | 9.139164431 | 2.33E-10 | 7.457342153 | 1.41E-07 |
| *HAUS7* | 1.077143968 | 0.000217 | 1.469187295 | 0.001673216 |
| ***HIST1H1A1*** | 5.848215734 | 1.80E-18 | 5.856905392 | 0.002878737 |
| *HSD17B10* | 1.031920999 | 9.67E-11 | 1.433723942 | 6.97E-05 |
| *IDH3G* | 1.001882884 | 4.29E-11 | 1.434234676 | 0.007311695 |
| *JADE3* | 1.079089438 | 3.51E-12 | 1.429530196 | 6.81E-06 |
| *NAA10* | 1.016446794 | 8.32E-09 | 1.420341345 | 0.000765417 |
| *NCAM2* | 1.175437195 | 0.005286634 | 1.103948768 | 2.24E-07 |
| *NDUFB11* | 1.171103978 | 1.43E-08 | 1.88856872 | 0.000684263 |
| ***PCDHGA31*** | 5.474057123 | 1.58E-122 | 4.801058573 | 3.10E-63 |
| *PCDHGA4* | 2.086201001 | 2.37E-11 | 1.512198469 | 0.016392048 |
| *PCDHGB1* | 1.081975754 | 0.001407812 | 1.525777875 | 0.021808739 |
| *PCDHGB2* | 1.746665418 | 9.95E-10 | 1.827544024 | 0.007254347 |
| *PCDHGB3* | 2.323839531 | 5.77E-15 | 1.873734051 | 0.002575344 |
| *PDHA1* | 1.093747283 | 3.29E-17 | 1.509521746 | 1.13E-19 |
| *RPL10* | 1.023326738 | 0.00025 | 1.397910368 | 0.024534554 |
| ***SCML21*** | 1.115859018 | 4.57E-12 | 1.650203502 | 1.49E-07 |
| ***SLC15A41*** | 2.640470731 | 6.30E-25 | 1.152395578 | 0.005698916 |
| ***SLC16A21*** | 1.178133567 | 1.31E-05 | 1.771954006 | 6.49E-10 |
| *SLC30A8* | 1.506637641 | 0.0002 | 1.238069979 | 0.00434561 |
| ***TMEM132D1*** | 9.931158475 | 2.67E-13 | 8.251489265 | 1.03E-08 |
| *TSPAN7* | 1.175352888 | 1.61E-07 | 2.135303521 | 5.38E-10 |
| **ZNF37A1** | 4.860032685 | 3.23E-87 | 4.325495167 | 4.14E-22 |
| **ZNF41** | 1.13161708 | 9.43E-08 | 1.149731459 | 0.000458778 |

<sup>1</sup> These genes are dysregulated throughout differentiation.

### L48R variant increases *H3-3B* gene expression and has increased protein deposition in 2D neuronal cell types

Because histones orchestrate chromatin compaction, their deposition in the nucleus affects both genome organization and gene expression. We posited that differential H3.3 deposition between L48R and control cells might underlie some of the gene expression changes we observed. We directly queried the altered expression of *H3-3A* and *H3-3B* in cells harboring the L48R variant. Notably, expression of *H3-3B* was significantly upregulated in NPCs (adjusted p-value = 1.58E-9), while *H3-3A* expression was not significantly affected (adjusted p-value = 0.99) (Fig. S3A). Using RT-qPCR, we validated this RNA-seq result, finding that *H3-3B* mRNA levels are upregulated in NPCs, and subsequently observed the same effect in FBNs (Fig. S3B). We observed no change in total levels of H3.3 protein between control and L48R NPCs (Fig. S3C). We next stained FBNs for H3.3 to understand the deposition of H3.3 in these post-mitotic cells and quantified the corrected total cellular fluorescence of H3.3/DAPI (Fig. S3D). We observed a significant increase in this metric in L48R cells compared to control, suggesting that H3.3 is hyper-deposited.

### L48R variant alters chromatin accessibility and the occupancy of transcription factors important for neuronal development in 2D models

To understand how increased H3.3 deposition affected global chromatin accessibility, we performed Assay for Transposase-Accessible Chromatin with sequencing (ATAC-seq) in two biological replicates of L48R NPCs and isogenic controls. L48R substantially opens chromatin compared to the control, specifically at the TSS (Fig. 3A, Fig. S3E) Using DiffBind^43^, we identified 350 differentially accessible sites (FDR ≤0.05, p ≤ 0.01), which were most commonly within 1kb of a promoter (40.86%) and in distal intergenic regions (21.43%) (Fig. S3E, Supplementary File 2). Of these 350 regions, 253 were significantly more open (log2fold-change ≥ 1) and 97 regions were significantly less open (log2fold-change ≤ −1) in the L48R NPCs compared to isogenic controls (Fig. 3B). We performed a GO analysis using all 350 sites and found that he most common terms were related to regulating neuronal development, axonogenesis, and cell growth (Fig. 3C). *PLXNA3*, a gene important for axon extension^52,53^, had significantly more open chromatin in the L48R NPCs compared to controls (Fig. 3D). *RBFOX1*, an RNA-binding protein that regulates neuronal splicing and is implicated in a range of NDDs^54^ had regions of more compact chromatin compared to control (Fig. S3H). *RBFOX1* expression is significantly decreased in NPC, validating the ATAC seq (Fig.S3I). Taken together, we found changes in H3.3 deposition in NPCs harboring the L48R variant. The resulting aberrant chromatin accessibility occurred at sites important for early neural development (*RBFOX1*) as well as later stages of axon and synapse development (*PLXNA3*), suggesting that some genomic regions may be improperly poised for transcription in cells that harbor the L48R variant.

**Figure 3.**
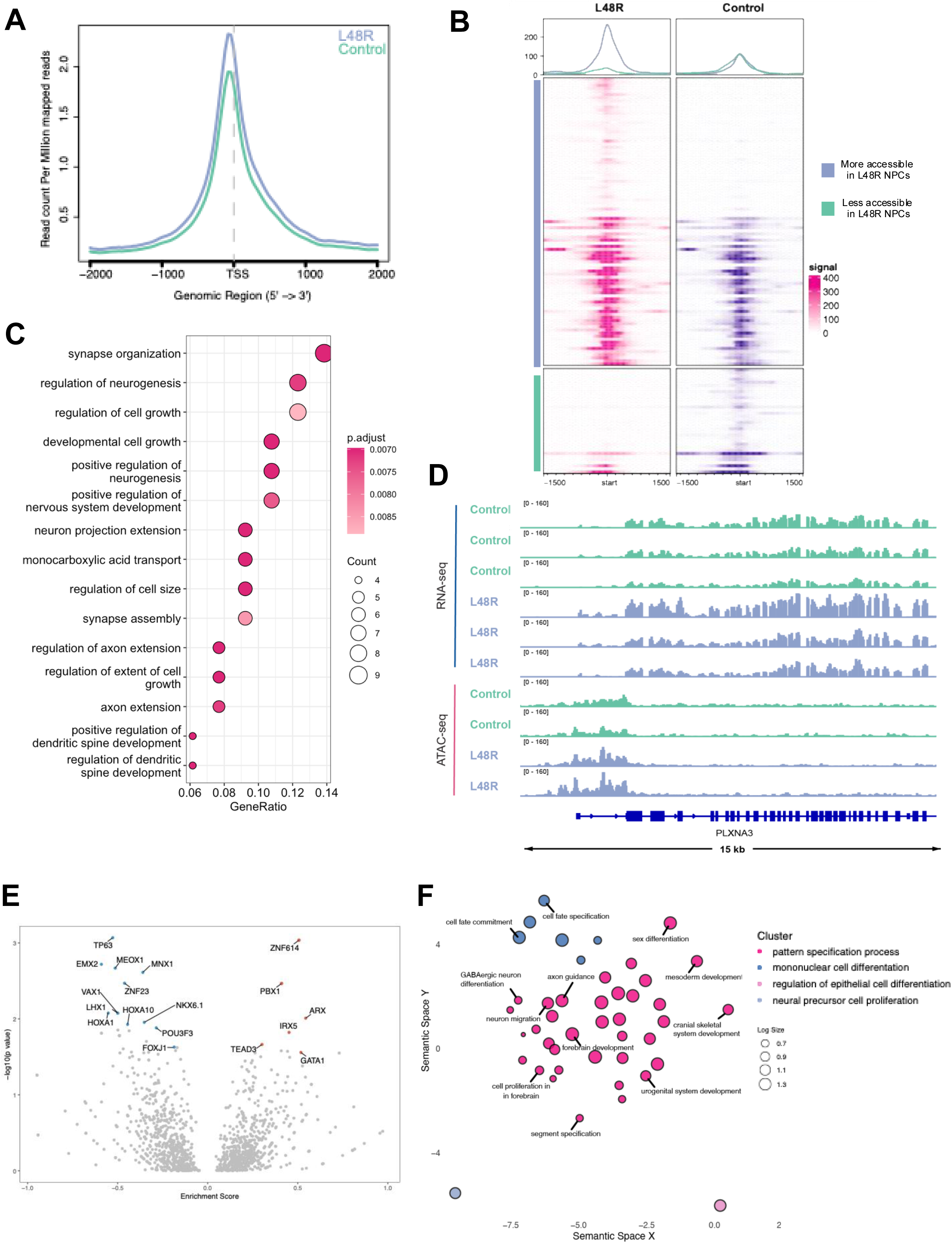
L48R variant alters chromatin accessibility and the occupancy of transcription factors important for neuronal development. A) Metaplot comparison of ATAC-seq average signal from control and L48R NPCs at all peaks, centered at the transcription start site (*n* = 2 Control, 2 L48R biological replicates). . B) Genomic heatmap of differentially accessible sites grouped by sites that are more accessible in L48R (top, purple), and those that are less accessible in L48R NPCs (bottom, green). C) GO enrichment of regions with significantly increased chromatin accessibility, annotated by nearest gene, in L48R NPCs compared to control. D) RNA-seq (top, blue) and ATAC-seq (bottom, pink) **reveals chromatin accessibility changes at loci regulating axon guidance (PLXNA3) in L48R NPCs.** E) **Transcription factor analysis indicates increased occupancy of PBX1 and decreased occupancy of TP63, suggesting dysregulation of differentiation and neurotransmission pathways.** F) REVIGO summary of GO terms associated with transcription factors in E) clustered by similar GO terms.

Next, we interrogated how changes in chromatin accessibility affected transcription factor (TF) occupancy. Using ATACseqTFEA^55^, we performed an enrichment analysis and identified 64 TFs predicted to have differential occupancy between L48R and control. This analysis generates enrichment scores; positive scores indicate that the motif(s) for a TF are enriched in L48R cells, and negative scores indicate that the motif(s) are depleted. Significant factors were called with an adjusted p-value ≤ 0.05. Of these factors, *ZNF614*, *PBX1,* and *ARX1* had increased occupancy in L48R, while *TP63* and *EMX2* had decreased occupancy in L48R (Fig. 3E). Increased occupancy of these TFs supports our previous data that transcription is altered, with a specific effect on neurotransmission. Meanwhile, the factors that were depleted in L48R support the transcriptomic data that the L48R variant impairs NPC differentiation.

We performed a GO analysis of all factors with changes in occupancy and performed a dimensional reduction of these terms using REVIGO^36^ (Fig. 3F). The terms clustered into four groups represented by the terms: pattern specification process (cluster 1), mononuclear cell differentiation (cluster 2), regulation of epithelial cell differentiation (cluster 3), and neural precursor cell proliferation (cluster 4) (Fig. 3F). Every cluster contains TFs important for fate specification in the developing nervous system. The most affected terms are important for transcription of genes required for neural precursor cell proliferation, stem cell differentiation and fate commitment, and forebrain development. To understand if altered TF occupancy might account for changes in gene expression, we used TFLink^56^ to find the targets of ZNF614, PBX1, and ARX (increased occupancy) as well as TP63 and EMX2 (decreased occupancy), and looked for changes in gene expression in our NPC RNA-seq dataset. The TFs with the greatest number of targets affected in this dataset were *PBX1* (12 up, 5 down) and *TP63* (32 up, 19 down) (Table 4) (Fig.S3G). *DPP6*, *GABRG3*, *GABRA5, CDH4, PCDHGA3, WNT7A*, and *PLXNA3*, genes with the largest differences in expression in L48R NPCs as compared to control, are all predicted targets of p63 regulation. While validation is required to show changes in binding/activity of these factors, these data support that the L48R variant affects TF binding by driving changes in chromatin accessibility, resulting in aberrant expression of genes required for NPC differentiation into mature neurons.

**Table 4.**
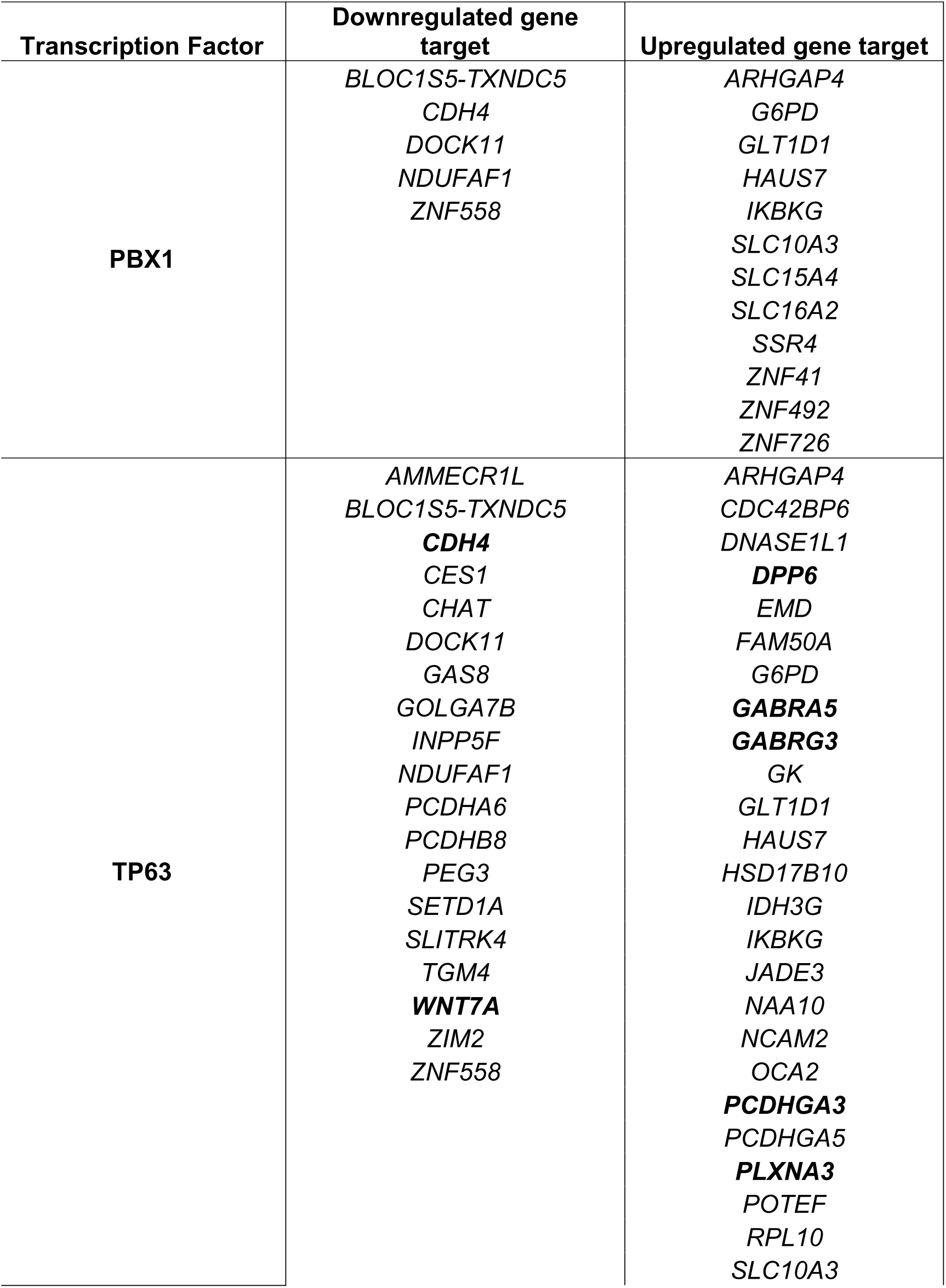
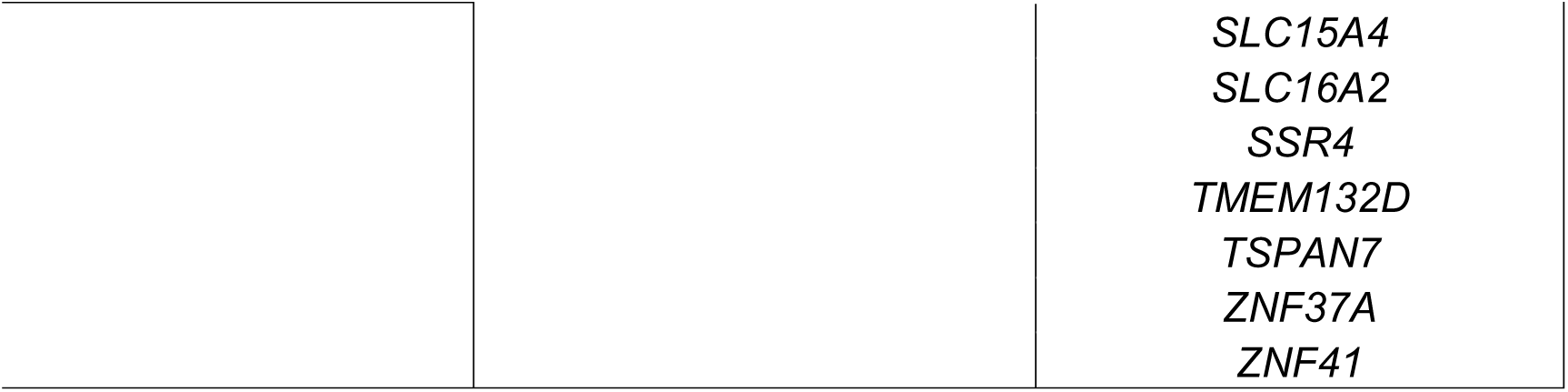
Targets of PBX1 (increased occupancy) and TP63 (decreased occupancy).

**Table 5.**
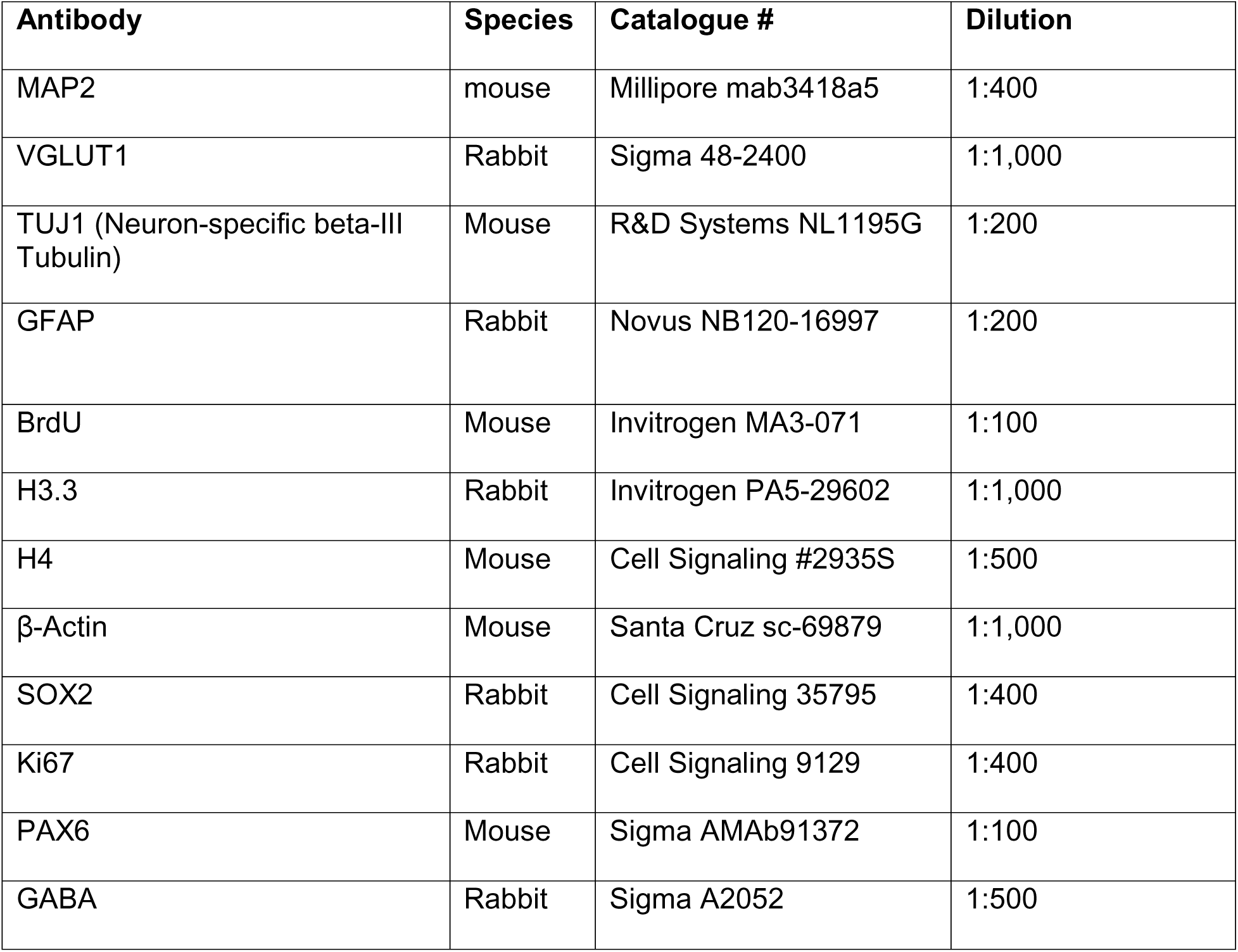
Primary antibodies used for protein-based experiments.

**Table 6.**
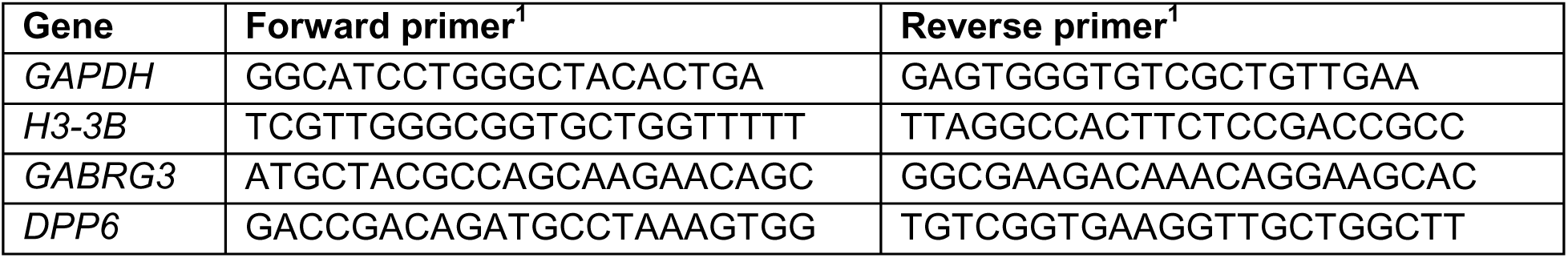
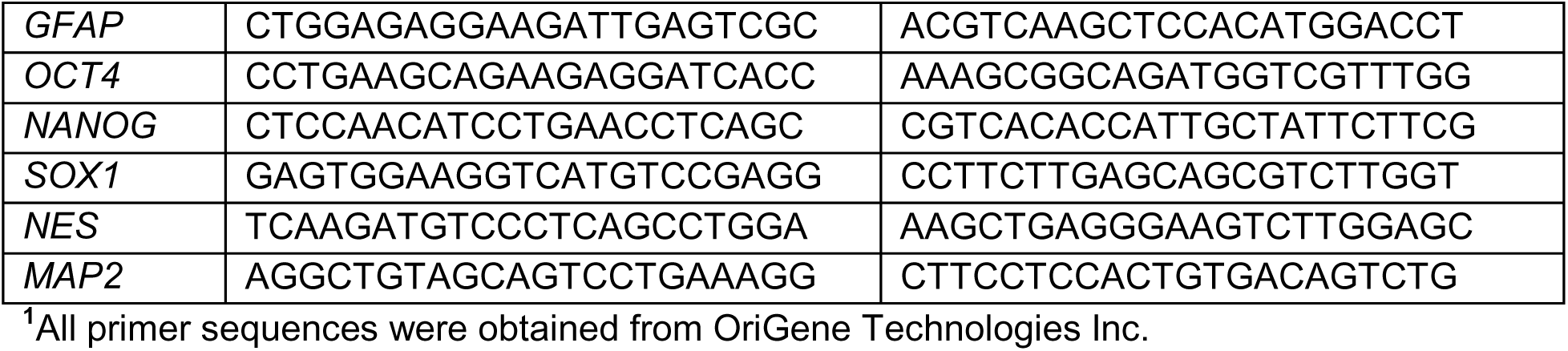
qPCR primers used to validate cell identity and validate RNA-seq results.

### Chromatin accessibility changes in L48R NPCs affect expression of genes important for cell adhesion and neurotransmission in 2D model

We next queried the overlap between ATAC-seq and RNA-seq datasets. First, we determined the relationship between changes in accessibility and gene expression by comparing the log2fold change in ATAC-seq reads binned by genes that were down, up, or unchanged in the RNA-seq. We found that downregulated genes had the greatest decrease in accessibility, while upregulated genes had the greatest increase in accessibility (Fig 4A). We next identified six genes that were down (designation from DESeq2 for RNA-seq and Diffbind for ATAC-seq) in both datasets (p ≤ 0.05, LFC ≤ −1) (Fig. 4B). *CDH4* was the most downregulated gene across these datasets (Fig. 4C, Table 1). We performed RT-qPCR gene expression analysis in iPSCs, NPCs, and FBNs to validate these results, and to understand *CDH4*’s expression during both development and maturation. In isogenic controls, *CDH4* expression increased as the iPSCs were differentiated to NPCs and FBNs, but when L48R iPSCs were differentiated to NPCs and FBNs, *CDH4* had nearly undetectable levels (Fig. 4D).

**Figure 4.**
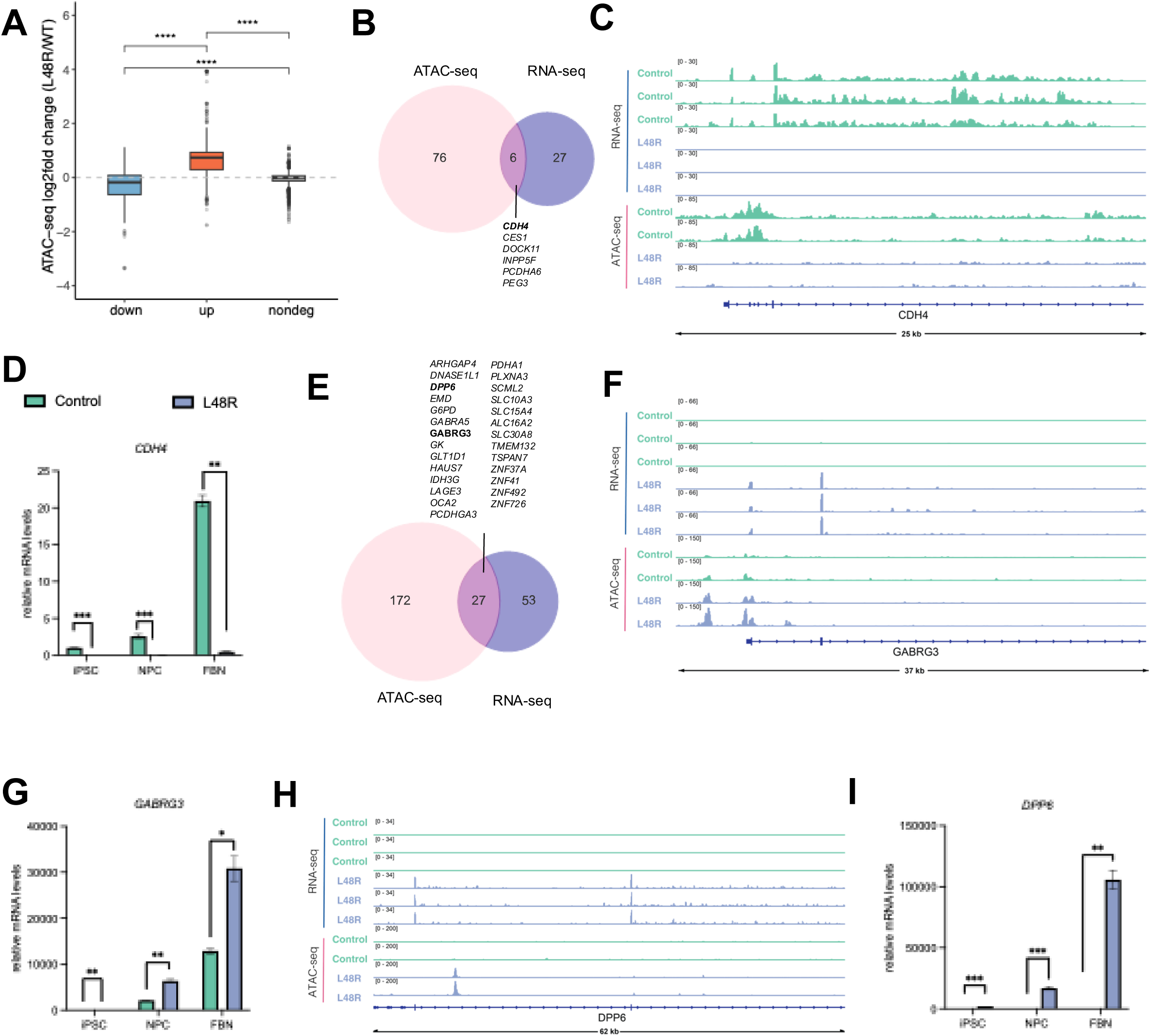
Chromatin accessibility changes in L48R 2D NPCs affect expression of genes important for cell adhesion and neurotransmission. A) ATAC-seq fold change (L48R/Control) for downregulated, upregulated, and unchanged genes in L48R NPC RNA-seq. Statistics were calculated using a t-test with Bonferroni correction. B-H) **Overlap analysis identifies upregulation of neurotransmission-related genes (e.g., GABRG3, DPP6) and downregulation of adhesion-related genes (e.g., CDH4) in L48R NPCs. These results suggest chromatin and transcriptional dysregulation contribute to impaired differentiation and neuronal maturation.**B) Venn diagram of overlap of significantly downregulated genes in RNA- and ATAC-seq. C) RNA-seq (top, blue) and ATAC-seq (bottom, pink) tracks for *CDH4*, a cell adhesion gene downregulated across all L48R cell types. Control tracks are shown in green, L48R in purple. D) RT-qPCR validation of *CDH4* mRNA levels from iPSC, NPC, to FBN. ΔΔ Ct values are normalized to *GAPDH*. p-values: iPSC = 9.834e-05, NPC = 0.0008733, FBN = 0.001522. E) Venn diagram of overlap of significantly upregulated genes in both datasets. F-G) RNA- and ATAC-seq tracks + RT-qPCR validation of *GABRG3*. p-values: iPSC = 0.002509, NPC = 0.008845, FBN = 0.02047. H-I) RNA- and ATAC-seq tracks + RT-qPCR validation of *DPP6.* p-values: iPSC = 0.0001059, NPC = 2.596e-05, FBN = 0.005359. Comparisons made via t-test and shown with brackets. *p ≤0.05, ** p ≤0.01, *** p ≤0.001

Then we identified 27 genes that were significantly upregulated in both datasets (p ≤ 0.05, LFC ≥ 1) (Fig. 4E). Notable genes include *GABRG3* and *DPP6*, two genes involved in neurotransmission (Table 1). *GABRG3* had more open chromatin at its promoter (Fig. 4F) and its expression increased as iPSCs were differentiated to NPCs and FBNs, with a significantly greater increase in L48R cells compared to control (Fig. 4G). A related gene, *GABRA5,* was also significantly up in both datasets (Fig. 4E, Fig. S3G). Further, DiffBind called eight different regions of *DPP6* more accessible in L48R NPCs: 1 exonic, 4 intronic, and 3 within 1kb of its promoter (Fig. 3H). When we analyzed the expression of *DPP6* in iPSCs, NPCs, and FBNs, we observed a 20,000-100,000-fold induction in L48R NPCs and FBNs (Fig. 3I). Taken together, our ATAC-seq and RNA-seq identified a set of genes important for neurotransmission, neuronal identity, and cell cycle progression with expression and chromatin architecture that are perturbed by the L48R variant in NPCs. The changes in *H3-3B* expression and H3.3 deposition, coupled with altered TF occupancy, suggest that L48R drives altered chromatin accessibility, resulting in abnormal gene expression and subsequently affecting NPC differentiation into neurons and glia.

### L48R variant affects radial glia and alters neuronal maturation in 3D dorsal forebrain organoids

Using an orthogonal model system to further analyze the neurodevelopmental consequences of the L48R variant, we utilized 3D dorsal forebrain organoids (DFBOs) to understand cortical development, neuronal maturation, and organization. DFBOs have been used in prior studies of neurodevelopmental disorders, including models of autism spectrum disorder showing altered excitatory/inhibitory balance and WNT signaling dysregulation (PMID: 26186191), as well as investigations into radial glia behavior and mitotic defects in Miller-Dieker syndrome, a severe lissencephaly (PMID: 28111201).

To study the effects of the L48R variant on early brain development, we converted isogenic control and L48R 2D iPSCs into 3D DFBOs^45^ and analyzed them at different time points (Fig. S4A). DFBOs were collected at day 25 (D25) to analyze the neuroepithelial expansion (NE) and at day 62 (D62) to analyze maturation. We analyzed organoid size across stages of development and observed that L48R organoids were larger at day 25 compared to isogenic controls, but this difference was not observable at later time points (Fig. 4A). L48R DFBOs at D62 had fewer NE buds at their edges than isogenic controls (Fig. S4B).

At D25 and D62, we observed autonomously organized radial glia in both sets of organoids by staining them with SOX2 (Fig. 5B,C). While there was no difference in SOX2 staining at D25 (Fig 5B), L48R DFBOs had significantly less expression of this marker at D62 (Fig. 5 C). Next, we stained organoids with CTIP2, a marker of intermediate progenitor neurons. While not significant, we observed that L48R DFBOs had more CTIP-positive cells at both time points (Fig. S4D). During neurodevelopment, radial glia undergo both symmetric division, allowing for self-renewal, and asymmetric division, which produce intermediate progenitor neurons^57^ further supporting that the pool of radial glia are being depleted through a skewing towards asymmetric division, leading to increased progenitors committed to a neuronal fate. L48R DFBOs had decreased TUJ1 staining at D25, but this difference was not present at D62 (Fig. 5B,C). This may suggest that, while increased CTIP2+ intermediate progenitor cells are available, they are less able to mature into TUJ1+ neurons within this D25 to D62 timeframe.

**Figure 5.**
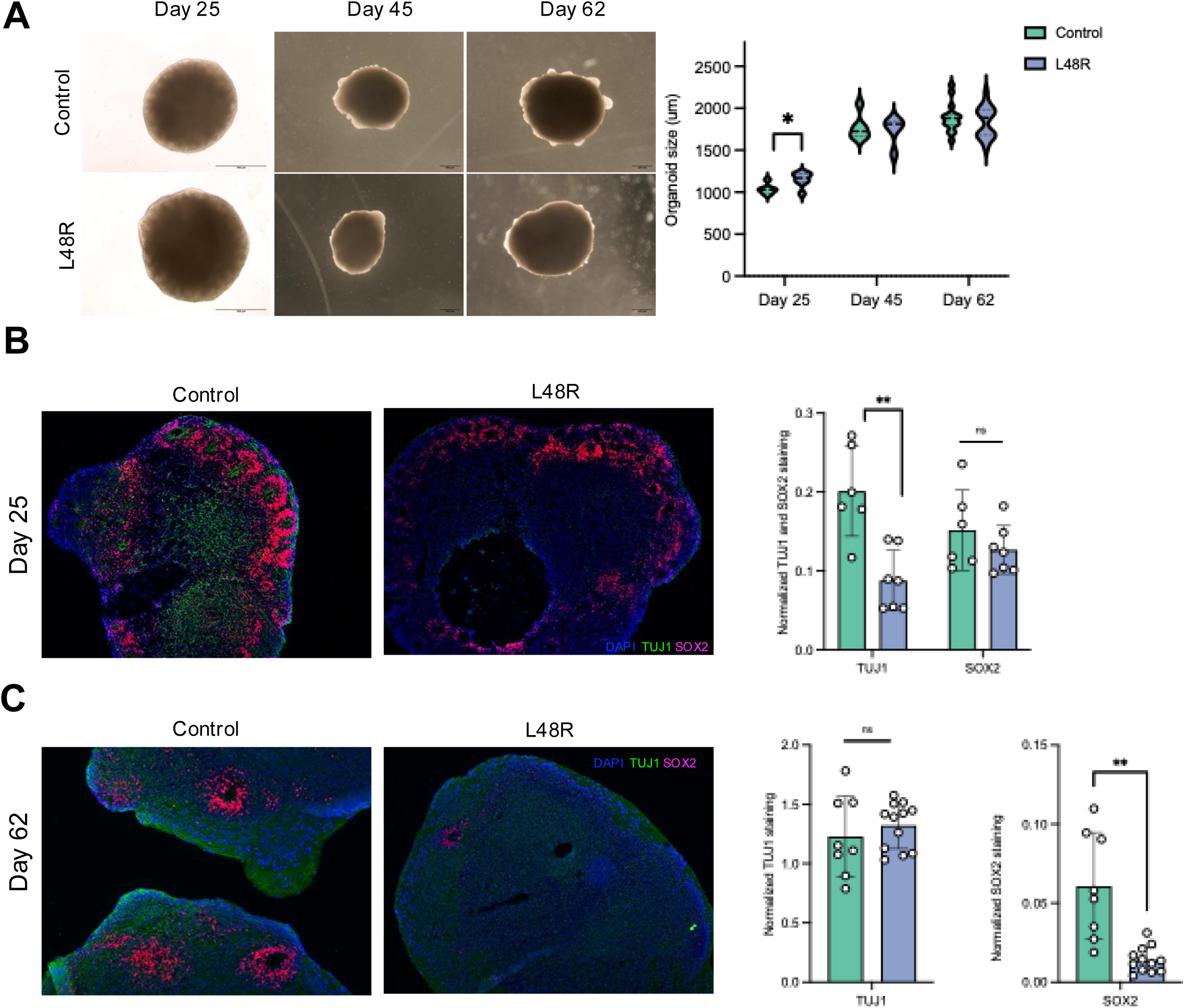
L48R variant perturbs progenitor populations in 3D DFBOs. A) Left: Representative images of organoids at D25, 45, and 62. D25images were taken at 10X magnification, D45 and D62 images were taken at 4x magnification, scale bar = 500μm Right: Quantification of organoid size using Qupath^67^. D25 n = 6 control, n = 7 L48R; D45 n = 6 control and L48R; D60 n = 15 control, n = 16 L48R. D25 p-value = 0.01684,. B) Organoids collected at D25 were stained with neuronal marker TUJ1 (green), progenitor marker SOX2 (pink) and DAPI (blue). **(C).** D25 n = 6 control, n = 7 L48R. TUJ1 p-value = 0.002887. C) Organoids collected at D62 were stained for the same markers as in B). Quantification of staining is shown on the right. D62 n = 8 control, n = 12 L48R SOX = 0.005652. Comparisons made via t-test and shown with brackets. *p ≤0.05, ** p ≤0.01, *** p ≤0.001

Since we observed dysregulation in the radial glia population, we stained D25 and D62 organoids with the cell proliferation marker Ki67. At D25, we observed a slight, but not significant, increase in Ki67-positive cells in L48R organoids. By D62, however, there was a significant decrease in Ki67-positive cells, indicating a loss of proliferative activity over time (Fig. 6A). To further probe cell cycle dynamics, we performed a BrdU incorporation assay, which labels cells in S phase. At day 25, BrdU incorporation was significantly increased in L48R organoids, suggesting early expansion of the proliferative progenitor pool. By day 62, however, BrdU incorporation was significantly decreased in L48R organoids compared to controls (Fig. 6B,C). Taken together, these data suggest that the L48R variant promotes early accumulation of progenitor-like cells that readily enter S phase (increased BrdU at D25), but do not show a sustained increase in overall cycling (non-significant Ki67 increase at D25). Over time, these progenitors appear to exit the cell cycle prematurely, as indicated by reduced Ki67 and BrdU labeling at D62, along with decreased SOX2 expression (Fig. 6D). These results suggest that L48R radial glia may have a functional deficit in maintaining self-renewing, symmetric divisions, leading instead to an early shift toward asymmetric divisions and the expansion of intermediate progenitor populations relative to control.

**Figure 6.**
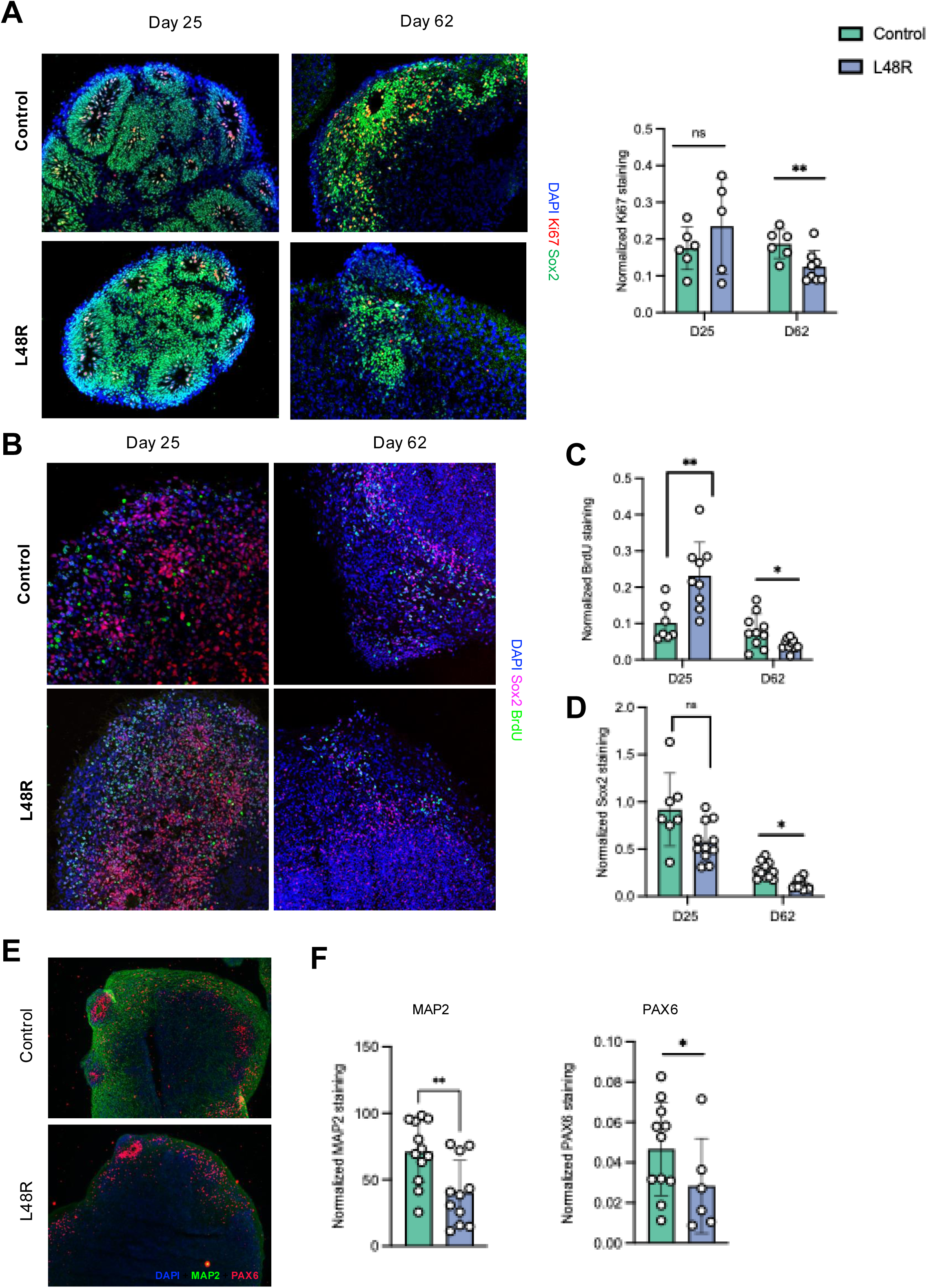
L48R DFBOs display abnormal proliferation and fewer mature neurons. A) Organoids stained for SOX2 (green) and proliferative marker Ki67 (red) at both day 25 and day 62. Images are quantified on the right. B-C) BrDU incorporation in control (top) and L48R (bottom) organoids at day 25 (left) and day 62 (right) with quantification (C). D) Quantification of SOX2, SOX2 p-values: D25 = 0.08127, D62 = 0.001318. E) D62 DFBOs stained with MAP2 (green) and PAX6 (red). R epresentative images taken at 10X are displayed **L48R DFBOs exhibit reduced mature neuronal marker MAP2.**. p-value = 0.006039. PAX6 staining is quantified in Fig S4C. Comparisons made via t-test and shown with brackets. *p ≤0.05, ** p ≤0.01, *** p ≤0.001

To understand how the L48R variant affects neuronal maturation, we stained organoids with mature neuronal marker MAP2, which stains the dendrites of mature post-mitotic neurons. We stained DFBOs collected at D62 with MAP2 and observed significantly lower levels in the L48R DFBOs (Fig. 6E,F). These results also validate our findings in 2D FBNs. We also observed a decrease in PAX6, another radial glia marker in these organoids (Fig 6E,F). To this point, we identified that L48R DFBOs had fewer proliferating radial glia, an expanded pool of intermediate progenitors, and fewer mature TUJ1+ neurons at D62. However, from these experiments alone, it is not clear whether these differences in cellular composition affected the function of these organoids.

To begin to answer this question, we looked at the expression of GABA, as our sequencing experiments elucidated an increased expression of GABA receptors *GABRG3* and *GABRA5*. We stained for GABA in D62 DFBOs and observed more GABA staining in the L48R condition (Fig. 7 B). Further, we observed significantly higher mRNA expression of *GABRG3* in L48R DFBOs (Fig. 7 C), validating our sequencing results in an orthogonal model. Taken together, our work in DFBOs indicates that the L48R variant affect both symmetric radial glial proliferation and asymmetric radial glia division, ultimately leading to defects in neuronal maturation. Further, these results suggest that the L48R mutation altered neuronal subtypes, potentially leading to an imbalance of excitatory/inhibitory signaling.

**Figure 7.**
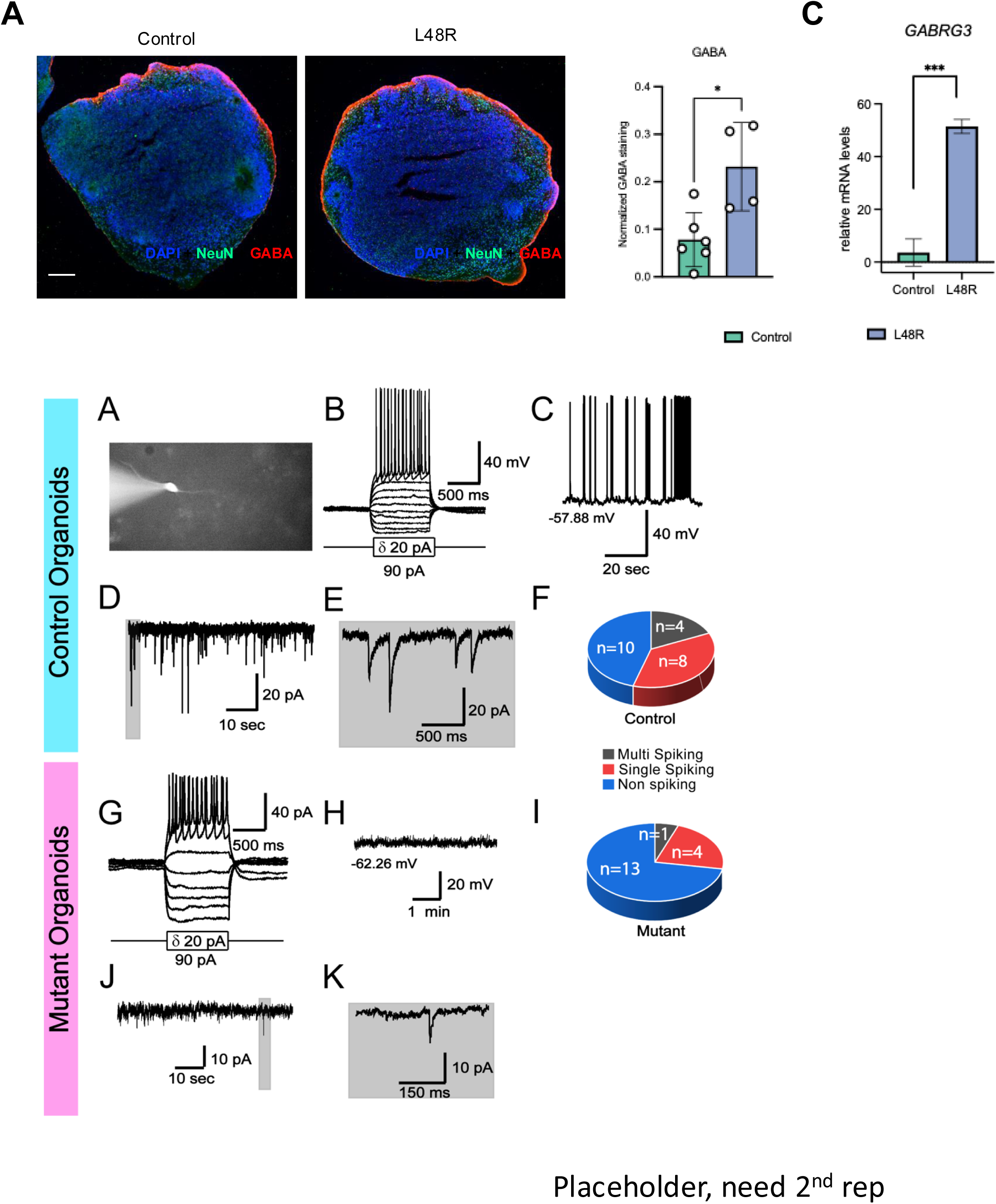
L48R DFBOs display increased GABA abundance. A-B) 10X images of D62 DFBOs stained with GABA (red) and NeuN (green). . p-value = 0.03561. C) RT-qPCR quantification of *GABRG3* levels in n = 4 DFBOs for both genotypes. p-value = 3.597e-05. D.a-j) Neurons in L48R DFBOs show less electrical activity than those in control DFBOs. **a.** Whole-cell patch clamp recording of a neuron from a control DFBO, showing action potential firing upon current injection (baseline membrane potential at - 70 mV). **b.** Percentages of neurons from the control organoids show various responses to current injections. **c.** Voltage-clamp recording trace showing spontaneous excitatory postsynaptic currents (sEPSCs) at a holding potential of −70 mV. **d.** Enlarged trace in the shaded area in panel C. **e.** Percentages of neurons with or without sEPSCs from the control organoids. **f.** Whole-cell patch clamp recording of a neuron from an L48R organoid, showing action potential firing upon current injection (baseline membrane potential at −70 mV). **g.** Percentages of neurons from the L48R DFBOs showing various responses to current injections. **h.** Voltage-clamp recording trace showing sEPSCs at a holding potential of −70 mV. **i.** Enlarged trace in the shaded area in panel H. **j.** Percentages of neurons with or without sEPSCs from the L48R DFBOs. E) L48R DFBO have less GFAP positive glial cells. 10X images of D90 DFBOs stained with GFAP (red) and MAP2 (green). Images were quantified as described in (A). p-value = 0.0003. Data is collected from two biological replicates and n=16 organoids from the two replicates.

### Neurons from L48R variant showed decreased electrical activity in 3D dorsal forebrain organoids

We next investigated the electrophysiological properties of neurons from control and L48R mutant 3D DFBOs via patch clamp recording (Fig. 7D). A significantly lower percentage of neurons in the L48R DFBOs fired action potentials in response to positive current injections compared to the control DFBOs (Fig. 7D a,b & f,g). Among the 45 neurons recorded from the L48R 3D DFBOs, 17.8% (8/45) were multi-spiking, 24.4% (11/45) single-spiking, and the remaining 57.8% (26/45) had no spiking (Fig. 7D e). In contrast, of the 48 neurons recorded from the control organoids, 37.5% (18/48) were multi-spiking, 37.5% (18/48) single-spiking, and the remaining 25.0% (12/48) neurons were not spiking (P<0.05, χ²-test) (Fig. 7D b).

Interestingly, for the multi-spiking neurons, neurons from L48R mutant DFBOs exhibited a higher firing frequency than those from control DFBO (Fig.S4E g) (Control: 12.1±1.4 Hz, n=18; Mutant: 18.0±0.9 Hz, n=8; P<0.05; student t-test). No significant differences were observed in resting membrane potential (RMP) or input resistance between neurons from control and mutant organoids (Fig. S4E e-i).

Furthermore, a subset of recorded neurons exhibited spontaneous excitatory postsynaptic currents (sEPSCs), reflecting synaptic inputs to these neurons. The percentage of neurons showing sEPSCs was significantly lower in the L48R mutant (8.9%; 4/45) compared to the control DFBOs (35.4%; 17/48) (P<0.05, χ²-test) (Fig. 7D e,j). No significant differences were found in the frequency or amplitude of sEPSCs (Fig. S4E h,i). These results suggest that the electrical activity in mutant DFBOs is comparatively less than that of their control counterparts. To investigate the glial population at day 90 of DFBO maturation, where we observed reduced electrical activity, we stained the DFBO with MAP2 and GFAP antibodies. We observed very low levels of glial cell expression in L48R organoids (Fig. 7E). These results indicate that the L48R mutation delayed neuronal maturation of DFBO by altering different cell types, further impacting their electrical activity.

## DISCUSSION

BLBS is a rare Mendelian syndrome with a mixed neurodevelopmental/neurodegenerative phenotype for which only palliative interventions are currently available. Prior work in patient-derived dermal fibroblasts, zebrafish, and mice have facilitated insights into the pathogenic mechanism of this neurogenetic disorder^5,7^. However, until now, the neurologic phenotype of BLBS had never been interrogated in a human model of neurodevelopment. In the first reported human iPSC-derived model of BLBS, we show that one BLBS causative variant, *H3-3B* p.L48R, impairs neuronal differentiation and maturation in both 2D iPSC and 3D organoid systems. We find that the identity of cells at the iPSC stage is largely unaffected by the L48R variant (Fig. 1B), but there are changes in cell cycle regulators and cell adhesion molecules that affect differentiation to NPCs (Fig. 2A, Fig. S2B). NPCs have decreased expression of fate markers like *NES* and *SOX1* (Fig. 1C), and factors important for differentiation including *CDH4, WNT7A* and *L1CAM* (Fig. 2B-D). Although we saw gene expression changes, total Forse1 and NCAD positive NPC population in the 2D FBN is not affected in our FACS experiments indicating subtle gene expression changes in iPSC are not affected overall NPC differentiation. Where as L48R FBNs express reduced MAP2 (Fig 1D, E) and have widespread transcriptional dysregulation, including decreased immediate early genes (Fig 2E) and increased astrocyte markers (Fig. S2D). These results indicate that L48R specifically disrupts NPCs, further impacting the neuronal differentiation and maturation. . Given the essential roles of these processes, such dysregulation could explain the near-universal neurologic features of BLBS. . Further, myelin-related anomalies in individuals affected by BLBS account for 46% of findings on MRIs^9^. Clinical evidence coupled with FBN transcriptomic data suggest a potential shift in progenitor cell fate towards glia, or a competitive disadvantage for neurons harboring the mutation, questions warranting further investigation. .

To further interrogate the drivers of these changes in gene expression in NPCs and FBNs harboring the L48R variant, we investigated how the expression of *H3-3B* itself was affected, finding that *H3-3B* expression was upregulated in both L48R NPCs and FBNs (Fig S3A,B). There was also increased H3.3 deposition in L48R FBNs (Fig. S3 D). This effect on its own gene expression suggests that it might be acting through a gain of function mechanism ^58^, meaning that gene disruption therapies could be a viable therapeutic option for individuals with this particular variant. Next, we defined the chromatin landscape of L48R NPCs using ATAC-seq and identified changes in chromatin accessibility at genomic regions important for axon guidance, neuron projection, and synapse organization (Fig. 3C, D). We also found changes in the occupancy of TFs that regulate processes like axon guidance and GABAergic neuron differentiation (Fig. 3E, F). Previous studies using iPSC-derived neural progenitor cells and neurons have demonstrated that chromatin accessibility dynamics are essential to proper neurodevelopment and are often disrupted in neurodevelopmental disorders such as autism spectrum disorder and epilepsy (PMID: 32732423; PMID: 39190448; PMID: 33545232). Our findings extend these observations by implicating a specific histone variant mutation in chromatin dysregulation, providing a direct molecular mechanism. Previous work done in embryonic stem cells showed that H3.3 promotes chromatin accessibility at promoters, and loss of H3.3 impairs transcription factor binding (PMID: 36782260). The fact that H3.3 L48R increases, rather than decreases, chromatin accessibility, further supports that this variant is gain of function.

While we were able to broadly define the chromatin landscape of L48R NPCs using ATAC-seq,, chromatin immunoprecipitation with sequencing (ChIP-seq) for H3.3 and the distribution of active/repressive hPTMs will provide more mechanistic insight underlying the changes in chromatin states. ChIP-seq specifically for L48R H3.3 would also indicate changes in chromatin accessibility and gene expression at the sites were due directly to altered H3.3 deposition.

When integrating our multi-omic data, we found that downregulated genes from RNA-seq had the greatest decrease in accessibility, while upregulated genes had the greatest increase in accessibility (Fig 4A). he RNA- and ATAC-seq datasets overlapped at a few key sites, and generally had more upregulated genes/open chromatin in L48R NPCs compared to isogenic controls (Fig. 4E). Many genes were related to key neurodevelopmental processes, such as cell adhesion and neurotransmission. The consistent downregulation of *CDH4*, a gene that encodes a cadherin fundamental for cell-cell interactions in the central nervous system, might be one of the first causes of differentiation deficits in L48R cells, preventing proper migration during neurodevelopment. The upregulation of *DPP6* and GABA receptors *GABRG3* and *GABRA5* suggest that L48R cells have impaired neurotransmission with decreased neuronal activity ^1^. This altered excitatory/inhibitory balance has been observed in several NDDs and is considered a unifying mechanism underlying this group of disorders.

Our work in 3D DFBOs recapitulate many of our findings in 2D cultures, and in what is seen clinically in affected individuals. When we measured the size of the DFBOs as they matured, we found that they were initially larger, and then normalized in size compared to control (Fig. 5A). At the time of last evaluation, the individual harboring the L48R variant in the BLBS cohort had microcephaly^5^. Interestingly, the growth phenotype of individuals with BLBS evolves over time. The first reported individual with BLBS presented with macrocephaly and overgrowth at birth but, at re-evaluation a decade later, exhibited microcephaly and severe undergrowth^5,8,62^. We found that this trend applies to other individuals affected by BLBS for whom follow-up data is available (https://doi.org/10.1016/j.xhgg.2025.100440). L48R DFBOs capture this phenotypic transition with increased organoid size and proliferation at D25, followed by reduced proliferation and progenitor cells at D62. Given that, reduced proliferation of neuronal progenitor cells is the basis for primary microcephaly (PMID: 30936766), we believe that this L48R DFBO model will be an effective model to study the mechanism driving this phenotype.

Another significant benefit of this DFBO model is that we could interrogate the impact of the L48R variant on multiple cell types at once. Our SOX2 and PAX6 staining show that there is a decrease in radial glia between D25 and D62 (Fig. 5B-C, Fig. S4C). The CTIP2 staining, which shows an increase in intermediate progenitor neurons during this same window (Fig. S4D), suggests that this decreased in radial glia is caused by a skewing towards the asymmetric division of radial glia, away from the expected homeostatic balance between self-renewing symmetric division and intermediate progenitor neuron-producing asymmetric division^57^. This is further supported by the Ki67, BrDU, and TUJ1 data, which show a decrease in proliferation accompanied by an increase in neurons between D25 and D62 (Fig. B-D). The decrease in radial glia may result from altered TF occupancy affecting pathways that regulate glial fate commitment and self-renewal. Thus, L48R cells ultimately differentiate into similar numbers of neurons at the cost of maintaining their progenitor identity. However, the D62 L48R neurons have delayed maturation (Fig. 6A), potentially because of dysregulated TF occupancy, or cell death caused by grossly dysregulated gene expression and chromatin accessibility. Further, our whole-cell patch clamp analysis indicated L48R DFBO has significantly less electrical activity and these organoids have very low glial cells (Fig. 7D, E). Taken together, these results indicate that L48R progenitors are less able to proliferate over the longer periods of organoid maturation and affected maturation of neurons and their electrical activity. The exhaustion of a replicative pool which only exists for a transient period during neurodevelopment might also explain why the central nervous system is universally affected but other organs are not.

We concluded our analyses at day 90 of maturation, where control DFBO started producing the low levels of GFAP positive glial cells, longer maturation periods are required to understand the significance of increased glial markers observed in the 2D FBNs, and their effects on glial production and proliferation in a 3D context will be informative, especially in understanding if these cells are skewed towards a non-neuronal fate. The use of cerebral organoids, which are less guided than DFBOs, would enable the study of additional cell types that make up hippocampal and retinal regions^64^. Oligodendrocyte-containing myelin organoids (myelinoids)^65^ will also be important models to understand the myelin-related anomalies observed in many individuals affected by BLBS.

Overall, our data suggest a model by which L48R acts through a gain-of-function mechanism leading to increased *H3-3B* expression that results in the hyper-deposition of H3.3 into the nucleosome, disrupting gene expression and chromatin accessibility. Functionally, this causes dysregulation of cell adhesion, neurotransmission, and the balance between excitatory and inhibitory signaling.

Alterations in GABAergic signaling, cell adhesion, and radial glia function are central to neurodevelopmental disorders (NDDs) and converge on core mechanisms of circuit formation. GABAergic dysfunction, a common feature in disorders like ASD and Rett syndrome, disrupts the excitatory/inhibitory balance essential for cortical function (PMID: 33772226, 21068835). Similarly, downregulation of adhesion molecules such as CDH4 can impair neuronal migration and synaptic organization, as seen in Fragile X syndrome (PMID: 39022311). Radial glia, critical for both progenitor expansion and neuronal scaffolding, are especially vulnerable to epigenetic and signaling perturbations, with defects linked to conditions such as microcephaly and lissencephaly (PMID: 28111201). Together, these observations underscore how epigenetic and transcriptional alterations like those caused by H3.3 L48R can converge on key developmental processes to yield circuit-level dysfunctions that underlie NDD pathogenesis.

The ability to define a proposed model with modifiable mRNA and protein level readouts, as well as functional consequences, establishes the necessary groundwork for future studies into therapeutic interventions. Specifically, our data show the upregulation of two GABA_A_ receptor encoding genes, *GABRG3* and *GABRA5,* in L48R NPCs. If GABA-mediated dysregulation is validated in future mechanistic work to underlie disease pathogenesis, this could be targeted through the repurposing of an FDA-approved class of GABA_A_ receptor selective positive allosteric modular antidepressants^66^. Further, because we found that *H3-3B* itself is upregulated, suggesting a not previously defined gain-of-function mechanism, targeting the mutated allele with an antisense oligonucleotide (ASO) or small interfering RNA may be a viable therapeutic. Finally, if future work identifies the dysregulation of hPTMs, perhaps an existing FDA-approved HDAC inhibitor could be repurposed for BLBS, which is being explored for MDEMs^2^. Our results from one model have highlighted a few therapeutic approaches, but intense future study into the mechanism of disease and the safety of possible interventions is needed to understand what therapeutics would be most beneficial for individuals affected by BLBS.

There are noteworthy strengths of our approach, which leverages 2D and 3D iPSC models coupled with sequencing and IF, but we acknowledge the limitations of this work. First, these studies were conducted in iPSCs that were derived from a single edited clone. To address this limitation, we performed at least two separate differentiations of iPSCs to each cell type queried; however, the gold standard would be to validate these findings in an independently generated cell line. Further, it is not clear if all findings from our XX (female) model would be recapitulated in an XY (male) model. Since there is an almost even number of males and females affected by BLBS^8^, these results would be strengthened if recapitulated in both an XX and XY model system. It is also important to note that 70 unique variants are reported to cause BLBS^8^, and we investigated the effects of only a single BLBS-causing variant. Because of the phenotypic variability observed in affected individuals and model systems^5,8,29^, future studies in isogenic cell lines from different origins harboring additional BLBS variants will be important in assessing the generalizability of these findings to have the greatest impact for affected individuals. In sum, we developed and interrogated a novel isogenic L48R iPSC line differentiated into 2D NPCs, 2D FBNs, and 3D DFBOs, uncovering critical insights into the pathogenesis of Bryant-Li-Bhoj syndrome (BLBS). These data support a model in which the L48R variant disrupts NPC proliferation and differentiation by altering chromatin accessibility at loci regulating axon guidance, cell adhesion, and synapse organization. These early developmental deficits prevent neuronal maturation and affect their electrical activity.

Furthermore, the dysregulation of GABA receptor pathways suggests a potential imbalance in excitatory/inhibitory signaling, with implications for neuronal network function. This study represents the first human iPSC-derived model of BLBS and highlights the critical role of chromatin dysregulation in neurodevelopmental disorders. Future work should focus on validating these findings in additional BLBS variants and assessing the therapeutic potential of antisense oligonucleotides or GABA receptor modulators. This model provides a foundation for understanding BLBS pathogenesis and advancing targeted therapeutic strategies for histone-associated neurodevelopmental syndromes.

## ACKNOWLEDGMENTS

We would like to sincerely thank the members of the BLBS community for sharing their information and journeys with us. Thank you also to Teodora Orendovici, PhD, Stephen Mahoney and Ashley Bushey from the CHOP High Throughput Sequencing Core for assistance with the ATAC-sequencing; Jean Ann Maguire, PhD from the CHOP iPSC core for generating the iPSC line; and Jonathan Billings, Kyra Booth, Tiancheng Wang and Kayleigh Ostberg from the CHOP Center for Applied Genomics for assistance with RNA-sequencing.

## CONTRIBUTIONS

**Data Collection**: RA, AKS, LMB, DLC, XMW, EAW, JPB, YW

**Data Analysis**: AKS, RA, ELD

**Figure Generation:** AKS, RA, EEL

**Conceived of study**: RA, AKS, EJB

**Drafted the manuscript**: AKS, RA, EEL

**Reviewed manuscript**: EEL, ELD, DLC, XMW, KJC, MM

## FUNDING

Support for this work was provided by NICHD F30 1F30HD112125 (EEL); NIGMS T32 5T32GM008638 (ELD); NHGRI T32HG009495 and the Eagles Autism Foundation (DELC); the Chan-Zuckerberg Initiative Neurodegeneration Challenge Network, as well as the Burroughs Wellcome Fund and the Hartwell Foundation (EJKB).

## DATA AVAILABILITY

The datasets generated during the current study will be available in the Gene Expression Omnibus repository.

## COMPETING INTERESTS

The authors declare no competing interests.

## Supplementary Figures

**Supplementary Figure 1.**
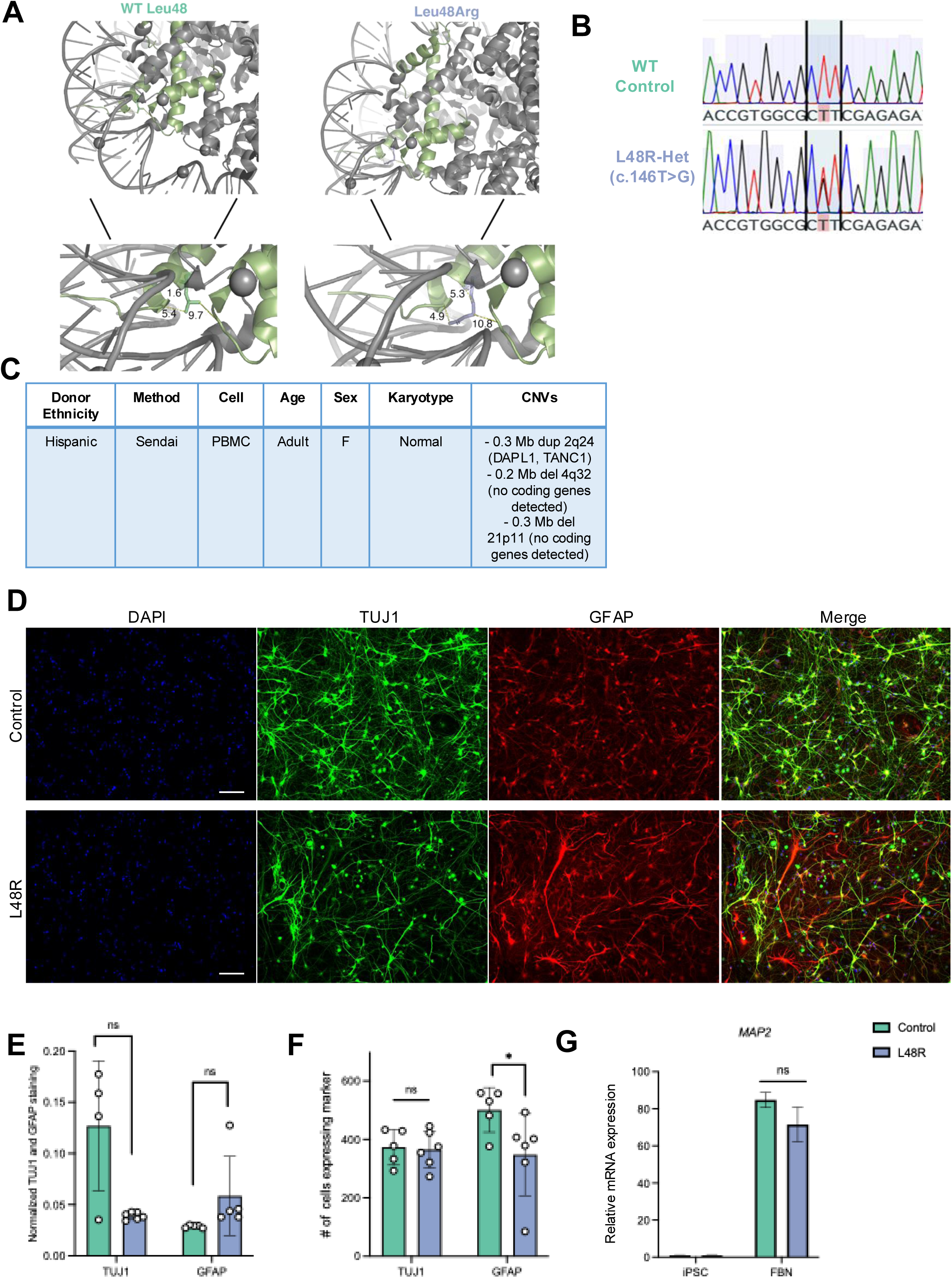
Characterization of 2D L48R iPSCs, NPCs and FBNs. A) 3D in silico structural model of the H3.3-containing nucleosome (PDB: 5X7X) with one of the two copies of H3.3 in purple; other histones in gray; and DNA in black/grey. The canonical Leu48 residue is shown on the left, the variant Arg48 is shown on the right in green. The distance in angstroms between H3.3 T45, H4 and DNA are displayed. B) Sanger sequencing verification of isogenic control (top) and heterozygous c.146T>C edit in CHOPWT14 cells (bottom). C) Description of CHOPWT14 iPSCs. Method = reprogramming method. D-G) Analysis of representative markers comparing iPSC and NPC differentiation using flow cytometry. D & F) Representative dot plots of flow cytometry results with antibodies specific to Forse1 and NCAD. At Day 0, iPSCs are devoid of Forse1 and NCAD, whereas by 20 days of NPC differentiation, cells are positive for both Forse1 and NCAD. E & G) Box plots showing the percentage of Forse1 and NCAD positive cells in four biological replicates. Samples were treated with secondary conjugated to Alexa 488 and data is represented as FITC signal against SSC-A. Data are analyzed from four biological replicates, each with a minimum of 10,000 cells analyzed for each condition. H) IF analysis in FBNs of TUJ1 (neurons) and GFAP (glia) collected at day 20. I) Quantification of n = 5 z-stacks for control, and n = 6 z-stacks for L48R FBNs. Corrected total fluorescence was calculated by dividing the fluorescence intensity of TUJ1 and GFAP by the fluorescence intensity of DAPI for each image. J) Quantification of the number of cells per image expressing each marker of interest. K) RT-qPCR analysis of *MAP2* mRNA levels in iPSCs and FBNs. p-values iPSCs = 0.4915, FBNs = 0.2222.

**Supplementary Figure 2.**
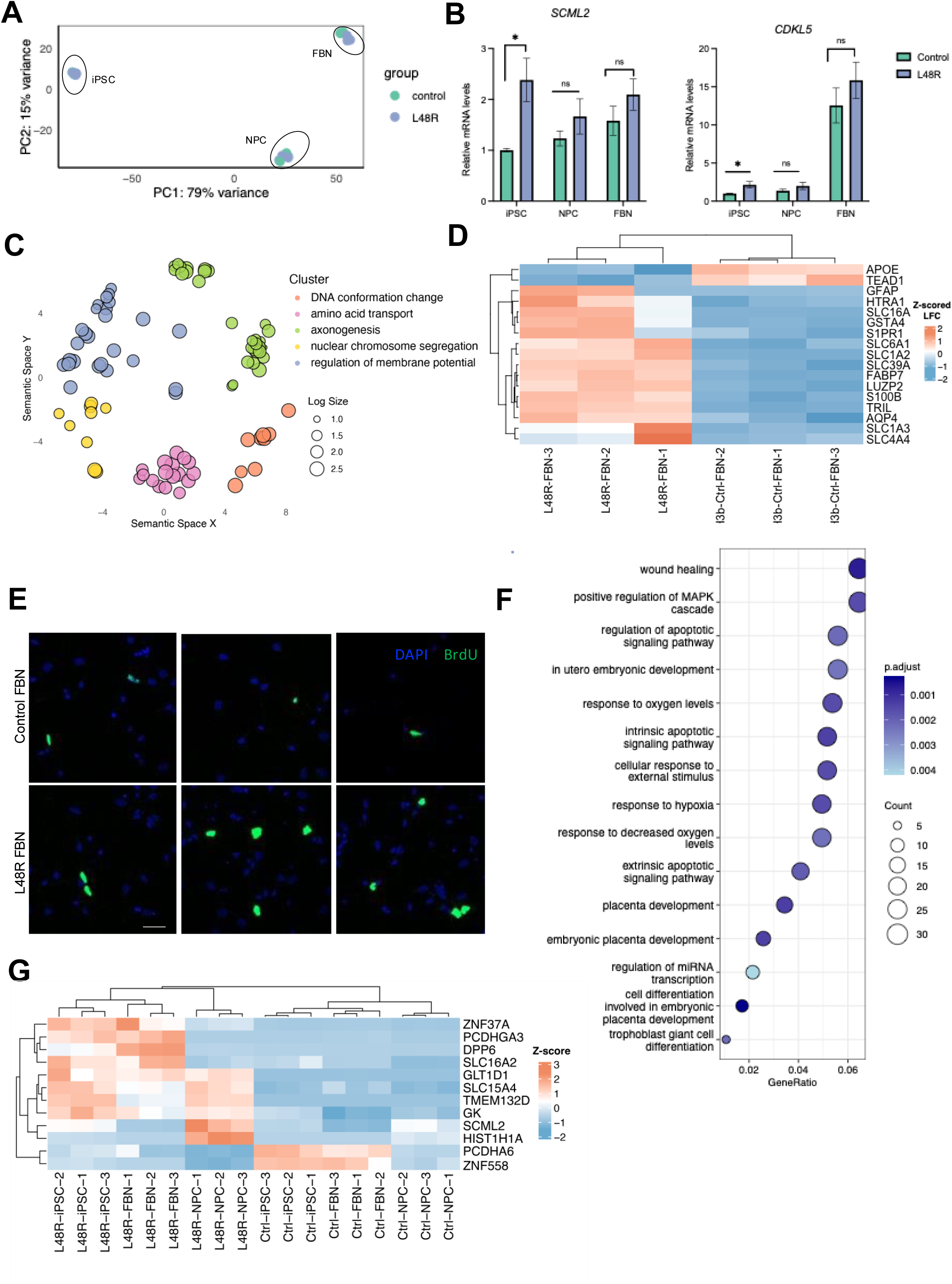
Transcriptomic dysregulation induced by L48R variant. A) PCA of L48R and control iPSCs, NPCs and FBNS, colored by genotype. B) RT-qPCR for SCML2 and CDKL5 to validate findings from bulk RNA-Seq. C) REVIGO summary of GO terms associated with significantly upregulated genes in FBNs, clustered by similarity. D) Heatmap of z-scored LFCs of astrocyte genes (from ^37^). E) BrdU labeling of control and L48R FBNs. F) GO plot of top 15 pathways downregulated in L48R FBNs compared to control FBNs. F) GO plot of significantly dysregulated genes in both L48R NPCs and FBNs compared to control.

**Supplementary Figure 3.**
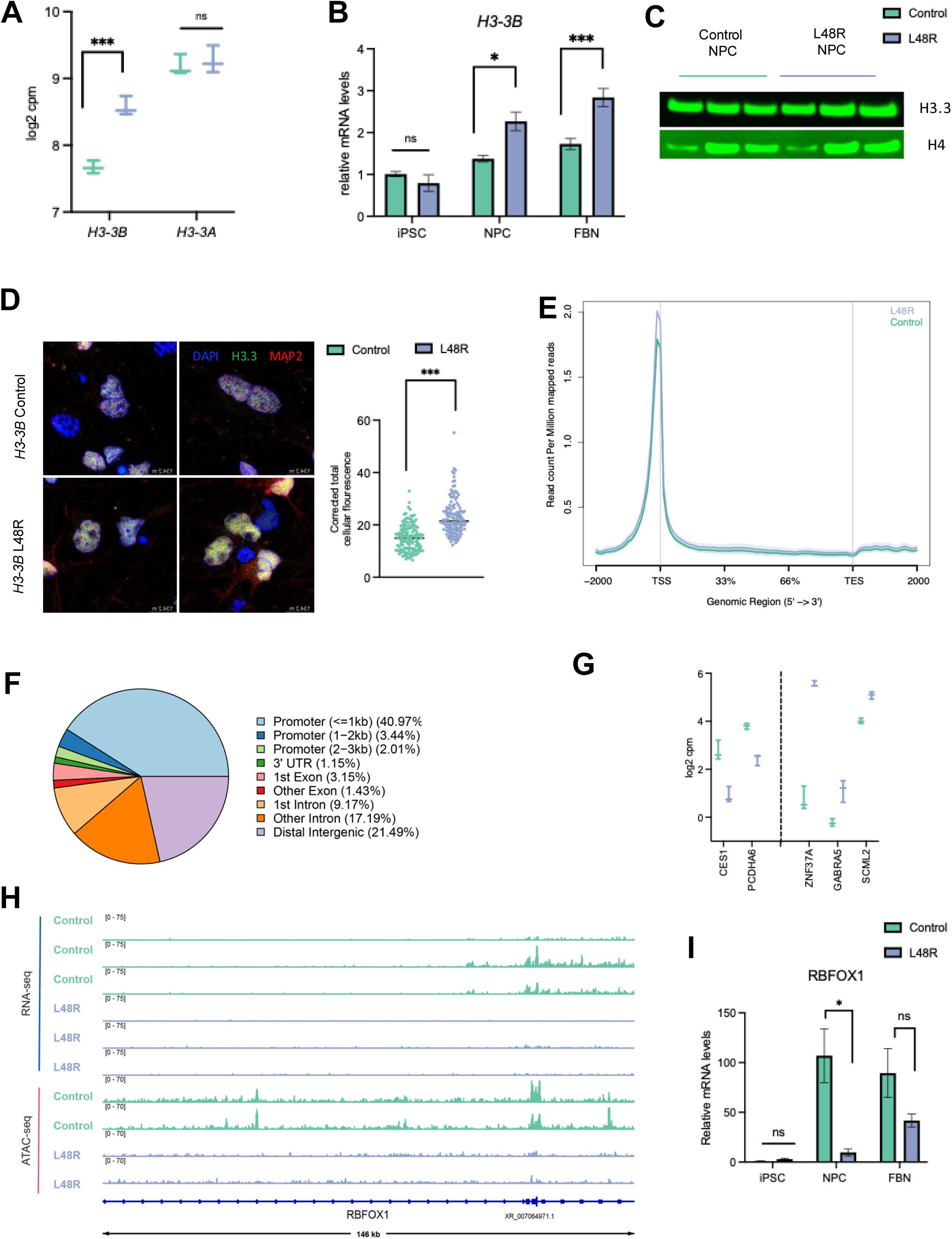
H3.3 deposition is altered in L48R cells, resulting in changes in chromatin landscape. A) log2 normalized counts of *H3-3B* and *H3-3B* from the NPC RNA-seq. B) RT-qPCR analysis of *H3-3B* in iPSCs (p-value = 0.3348), NPCs (p-value = 0.04321), and FBNs (p-value = 0.0006555). B) RT-qPCR validation of *H3-3B* expression in iPSCs, NPCs and FBNs. p = 0.051 in iPSCs, 0.0015 NPCs, p=0.4472 in FBNS. C) Western blot on nuclear extracted H3.3 and H4 as a loading control. C) Western blot probing for H3.3 in cytoplasmic and chromatin-bound fractions. D) IF for H3.3 and MAP2 in Control and L48R FBNs (left). Quantification of H3.3 fluorescence (right). p-value < 2.2e-16. E) Metaplot comparison of ATAC-seq average signal from control and L48R NPCs at all peaks, showing the region spanning +/- 2kb from the TSS (*n* = 2 control, 2 L48R biological replicates. F) Pie plot of the genomic annotations of sites that are differentially accessible in L48R vs control NPCs (identified using DiffBind) G) log2-normalized counts for selection of genes that were affected in both ATAC- and RNA-seq. Genes to the left of the dashed line were down and those to the right were upregulated. H) RNA- and ATAC-seq tracks for *RBFOX1*. I) RT-qPCR validation of gene expression changes in iPSCs, NPCs and FBNs. p-values: iPSC, NPC = 0.0153, FBN = 0.1113.

**Supplementary Figure 4.**
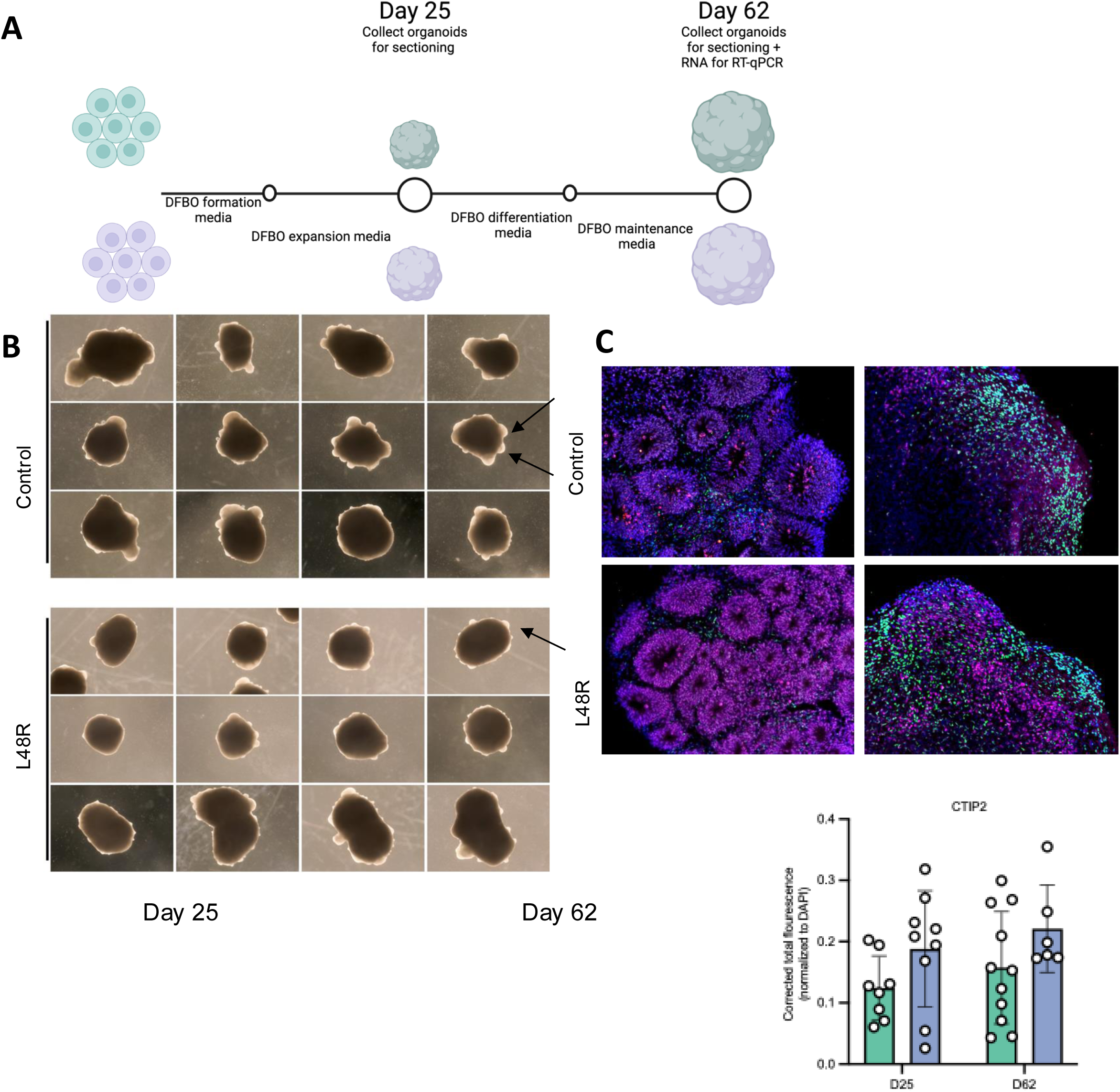
Cell identity analysis in 3D DFBOs corroborates 2D findings. A) Schematic of the dorsal forebrain organoid protocol employed in this study. Organoids were collected for sectioning at D25, and for RT-qPCR analysis and sectioning at D62. B) Images taken at 4X magnification of control (top) and L48R (bottom) organoids at D62 showing differences in NE buds (denoted by errors). C) Quantification of IF for PAX6 (red) at D62 (top) shown in Fig 6E. p-value = 0.3291 D) CTIP2 staining at D25 and D62, along with corrected total fluorescence quantifications of n = 6 organoids (control) or n = 4 organoids (L48R). E) Quantification of CTIP2 shown in D. D25 p-value = 0.1045, D62 p-value = 0.138. E. a-i) Intrinsic electrophysiological properties of neurons in control and L48R DFBOs. **a.** neuron from a control **(a)** and an L48R DFBO **(b)** filled with Alexa Fluor 488, depicting a bright soma with processes. **c, d.** Whole-cell patch clamp recording of neurons showing spontaneous activity from control and L48R DFBOs in current-clamp mode without current injection. **e-i**. Bar graphs showing mean (+SEM) and scatter plots comparison of resting membrane potential (**e**), input resistance (**f**), maximum spike frequency during current injection (**g**), sEPSC amplitude **(h)**, and sEPSC frequency **(i)** between control and L48R DFBOs. (\**P<* 0.05, ^NS^ *P* > 0.05; student t test). Data is collected from two biological replicates, 45 neurons from Control and 45 neutrons from L48R are analyzed.

## Supplementary Files

**Supplementary File 1**. DESeq analysis of RNA-seq experiments in iPSCs, NPCs, and FBNs.

**Supplementary File 2**. DiffBind results with significant changes in chromatin accessibility from the ATAC-seq in NPCs. Genomic regions were annotated with ChIPSeeker.

## Notes

### Competing Interest Statement

The authors have declared no competing interest.

### Summary of Updates

The revised version has additional experiments including flow cytometric analysis and patch clamp electrophysiologic analysis.

