## Supplemental Table for "A novel iPSC model of Bryant-Li-Bhoj neurodevelopmental/neurodegenerative syndrome demonstrates the role of histone H3.3 in chromatin dynamics, neuronal differentiation, and maturation"

### Table 1 Function and disease associations of the dysregulated genes in L48R compared to isogenic controls.

| Category | Data set | Gene name | Function | Disease Associations | References |
|---|---|---|---|---|---|
| Cell cycle progression / Differentiation | iPSC RNA-seq | *CDKL5* | Encodes a cell cycle inhibitor that promotes the differentiation of neuronal cells | Developmental and Epileptic Encephalopathy 2 | 47 , OMIM #300672 |
| | iPSC RNA-seq/RNA-seq + ATAC-seq overlap | *SCML2* | Encodes polycomb group protein that regulates the cell cycle by controlling the progression of cells from G1 into S phase | - | 48 |
| Cell adhesion/Cell migration | NPC RNA-seq + ATAC-seq overlap | *CDH4* | Encodes a classic cadherin, R-cadherin, a calcium dependent cell adhesion protein. Classic cadherins are important for tissue architecture across organs and specifically in the brain for neurite interactions | - | 68,69 |
| | NPC RNA-seq | *L1CAM* | Encodes L1CAM, which acts statically as glue between cells and promotes cell migration during neurodevelopment | X-linked hydrocephalus, Masa Syndrome | 70, OMIM #s 30335 307000 |
| | NPC RNA-seq | *FLNA* | Encodes Filamin A, an actin binding protein. It is required for neuronal migration to the cerebral cortex. This gene is also important for cardiac morphogenesis. | Periventricular Heterotopia, Otopalatodigital syndrome, Frontometaphyseal dysplasia, Melnick-Needles syndrome | 71,72 |
| | NPC-RNA-seq | *PCDHGB1, PCDHGB3, PCDHGA4, PCDHGA5, PCDHGB2, PCDHGA3, PCDHGC3, PCDHA6, PCDHB11* | Encode members of the clustered protocadherins, which are a family of cell-surface homophilic proteins. Protocadherins regulate axonal tiling and confer identity to neurons, which is required for the distinguishment between self and non-self. | ASD, schizophrenia | 49,51 |
| Axon guidance | RNA-seq + ATAC-seq | *PLXNA3* | Encodes the most widely expressed plexin receptor in fetal brain, plexin-A3[41], a class 3 semaphorin receptor known to be important for axon pathfinding in the developing nervous system and functions directly in signaling repulsion[42] | X-linked intellectual disability syndrome | 52 |
| | NPC RNA-seq | *WNT7A* | Required for neural stem/progenitor cell proliferation and self-renewal and adult mouse brains, stimulates the formation and function of excitatory synapses, increases density of dendritic spines | Fuhrmann syndrome, ASD | 73–75, OMIM#2289 |
| | NPC RNA-seq | *ARHGAP4* | Encodes a Rho GTPase that negatively regulates cell and axon motility | - | 76 |

| | | | | | |
|---|---|---|---|---|---|
| | NPC RNA-seq | *SEMA5A* | Encodes semaphorin-5A, a negative regulator of excitatory synapse formation | ASD | 77 |
| **Neuronal transcription factors** | ATAC-seq TF analysis | *PBX1* | Controls organ patterning during embryogenesis. In the brain, it regulates the survival of midbrain dopaminergic neurons and is an early regulator of subventricular zone neurogenesis | Parkinson's disease, CAKUTHED | 78,79, OMIM #6176 |
| | | *ARX* | Specifies forebrain patterning and growth, and specifically controls the development of telencephalic GABAergic neurons | Epilepsy and NDDs with or without structural defects | 80,81 |
| | | *TP63* | Encodes p63, a protein that works cooperatively with p53 to promote apoptosis during neuronal development, which is important for establishing neuronal connectivity | Ectrodactyly, ectodermal dysplasia, and cleft lip/palate syndrome 3 | 82, OMIM #604292 |
| | | *EMX2* | Expressed primarily in cortical progenitors where it regulates multiple features of cortical development including arealization and specifying positional identity of cortical cells | Schizencephaly | 83, OMIM # 26916 |
| **Neurotransmission** | RNA-seq + ATAC-seq | *DPP6* | *K*nown canonically as an auxiliary subunit of Kv4-mediated A-type $K^+$ channels, which act to delay excitation and regulate firing frequency. *DPP6* enhances the surface expression of these channels and is thought to accelerate their kinetics. It is also important for dendritic branching and synaptic development during neuronal development. | Schizophrenia | 84–86, OMIM #6000 |
| | | *GABRG3* | Encodes a $GABA_A$ receptor- γ3, which binds GABA, the major inhibitory neurotransmitter in the human brain. GABA regulates every step of neurogenesis and regulates NPC proliferation and migration through its -A receptors. | Epilepsy, ASD | 87–90 |
| | | *GABRA5* | encodes $GABA_A$ receptor subunit | Early onset epilepsy and developmental delay | 91 |
| **Immediate early genes** | FBN RNA-seq | *EGR1, FOSL1, JUNB, NPAS4* | Immediate early genes are rapidly upregulated in neurons in response to a wide variety of cellular stimuli and are important for synaptic plasticity. They provide the molecular framework for both a rapid and dynamic response to neuronal activity and lasting gene expression changes. | Anxiety, depression, schizophrenia | 50,92 |

**Table 2 Genes dysregulated across 2D iPSCs, NPCs and FBNs.**

| | iPSC | | NPC | | FBN | |
|---|---|---|---|---|---|---|
| **Gene ID** | **log2fold change** | **adjusted p-value** | **log2fold change** | **adjusted p-value** | **log2fold change** | **adjusted p-value** |
| *PCDHA6* | -2.193450727 | 2.17E-06 | -1.355795213 | 8.71E-09 | -1.308921624 | 0.045580571 |
| *ZNF558* | -1.148059517 | 0.004709638 | -1.128578936 | 8.37E-10 | -1.199734433 | 8.80E-05 |
| *GK* | 1.210861399 | 7.77E-10 | 1.030926781 | 8.37E-10 | 1.779059583 | 1.73E-09 |
| *SCML2* | 1.042489231 | 0.000268 | 1.115859018 | 4.57E-12 | 1.650203502 | 1.49E-07 |
| *SLC16A2* | 1.001951462 | 0.000177 | 1.178133567 | 1.31E-05 | 1.771954006 | 6.49E-10 |
| *SLC15A4* | 3.032307096 | 6.79E-27 | 2.640470731 | 6.30E-25 | 1.152395578 | 0.005698916 |
| *DPP6* | 3.988997109 | 4.29E-09 | 4.408206465 | 4.91E-65 | 6.226215688 | 1.75E-101 |
| *ZNF37A* | 5.871603628 | 9.76E-38 | 4.860032685 | 3.23E-87 | 4.325495167 | 4.14E-22 |
| *PCDHGA3* | 3.097728204 | 1.15E-05 | 5.474057123 | 1.58E-122 | 4.801058573 | 3.10E-63 |
| *HIST1H1A* | 2.471946327 | 8.11E-36 | 5.848215734 | 1.80E-18 | 5.856905392 | 0.002878737 |
| *GLT1D1* | 8.915844422 | 1.64E-09 | 9.139164431 | 2.33E-10 | 7.457342153 | 1.41E-07 |
| *TMEM132D* | 9.629365976 | 1.03E-11 | 9.931158475 | 2.67E-13 | 8.251489265 | 1.03E-08 |

**Table 3** Genes dysregulated in both L48R NPCs and FBNs compared to isogenic controls.

| | NPC | | FBN | |
|---|---|---|---|---|
| **Gene ID** | **log2fold change** | **adjusted p-value** | **log2fold change** | **adjusted p-value** |
| *ANKRD20A1* | -1.460915202 | 2.27E-06 | -1.439994876 | 4.64E-05 |
| *CDH4* | -3.590286774 | 2.77E-31 | -4.668488495 | 5.01E-62 |
| *PCDHA6* | -1.355795213 | 8.71E-09 | -1.308921624 | 0.045580571 |
| *PCDHB8* | -1.391417915 | 0.001425716 | -1.546283192 | 0.005391523 |
| *PEG3* | -1.908978559 | 1.35E-20 | -2.286904202 | 6.99E-06 |
| *TGM4* | -2.716613511 | 0.008999827 | -4.179492072 | 0.000408431 |
| *ZIM2* | -2.738228127 | 0.006096512 | -3.135289722 | 8.30E-07 |
| *ZNF558* | -1.128578936 | 8.37E-10 | -1.199734433 | 8.80E-05 |
| *DNASE1L1* | 1.186058534 | 0.001772585 | 1.378797988 | 0.005401369 |
| *DPP6* | 4.408206465 | 4.91E-65 | 6.226215688 | 1.75E-101 |
| *FAM58A* | 1.12636598 | 3.36E-12 | 1.207565785 | 0.013980524 |
| *GABRG3* | 4.22607955 | 9.14E-41 | 1.725418457 | 1.08E-07 |
| *GK* | 1.030926781 | 8.37E-10 | 1.779059583 | 1.73E-09 |
| *GLT1D1* | 9.139164431 | 2.33E-10 | 7.457342153 | 1.41E-07 |
| *HAUS7* | 1.077143968 | 0.000217 | 1.469187295 | 0.001673216 |
| *HIST1H1A* | 5.848215734 | 1.80E-18 | 5.856905392 | 0.002878737 |
| *HSD17B10* | 1.031920999 | 9.67E-11 | 1.433723942 | 6.97E-05 |
| *IDH3G* | 1.001882884 | 4.29E-11 | 1.434234676 | 0.007311695 |
| *JADE3* | 1.079089438 | 3.51E-12 | 1.429530196 | 6.81E-06 |
| *NAA10* | 1.016446794 | 8.32E-09 | 1.420341345 | 0.000765417 |
| *NCAM2* | 1.175437195 | 0.005286634 | 1.103948768 | 2.24E-07 |
| *NDUFB11* | 1.171103978 | 1.43E-08 | 1.88856872 | 0.000684263 |
| *PCDHGA3* | 5.474057123 | 1.58E-122 | 4.801058573 | 3.10E-63 |
| *PCDHGA4* | 2.086201001 | 2.37E-11 | 1.512198469 | 0.016392048 |
| *PCDHGB1* | 1.081975754 | 0.001407812 | 1.525777875 | 0.021808739 |
| *PCDHGB2* | 1.746665418 | 9.95E-10 | 1.827544024 | 0.007254347 |
| *PCDHGB3* | 2.323839531 | 5.77E-15 | 1.873734051 | 0.002575344 |
| *PDHA1* | 1.093747283 | 3.29E-17 | 1.509521746 | 1.13E-19 |
| *RPL10* | 1.023326738 | 0.00025 | 1.397910368 | 0.024534554 |
| *SCML2* | 1.115859018 | 4.57E-12 | 1.650203502 | 1.49E-07 |
| *SLC15A4* | 2.640470731 | 6.30E-25 | 1.152395578 | 0.005698916 |
| *SLC16A2* | 1.178133567 | 1.31E-05 | 1.771954006 | 6.49E-10 |
| *SLC30A8* | 1.506637641 | 0.0002 | 1.238069979 | 0.00434561 |
| *TMEM132D* | 9.931158475 | 2.67E-13 | 8.251489265 | 1.03E-08 |
| *TSPAN7* | 1.175352888 | 1.61E-07 | 2.135303521 | 5.38E-10 |
| *ZNF37A* | 4.860032685 | 3.23E-87 | 4.325495167 | 4.14E-22 |
| *ZNF41* | 1.13161708 | 9.43E-08 | 1.149731459 | 0.000458778 |

**Table 4 Targets of PBX1 (increased occupancy) and TP63 (decreased occupancy).**

| Transcription Factor | Downregulated gene target | Upregulated gene target |
|---|---|---|
| **PBX1** | *BLOC1S5-TXNDC5* | *ARHGAP4* |
| | *CDH4* | *G6PD* |
| | *DOCK11* | *GLT1D1* |
| | *NDUFAF1* | *HAUS7* |
| | *ZNF558* | *IKBKG* |
| | | *SLC10A3* |
| | | *SLC15A4* |
| | | *SLC16A2* |
| | | *SSR4* |
| | | *ZNF41* |
| | | *ZNF492* |
| | | *ZNF726* |
| **TP63** | *AMMECR1L* | *ARHGAP4* |
| | *BLOC1S5-TXNDC5* | *CDC42BP6* |
| | ***CDH4*** | *DNASE1L1* |
| | *CES1* | ***DPP6*** |
| | *CHAT* | *EMD* |
| | *DOCK11* | *FAM50A* |
| | *GAS8* | *G6PD* |
| | *GOLGA7B* | ***GABRA5*** |
| | *INPP5F* | ***GABRG3*** |
| | *NDUFAF1* | *GK* |
| | *PCDHA6* | *GLT1D1* |
| | *PCDHB8* | *HAUS7* |
| | *PEG3* | *HSD17B10* |
| | *SETD1A* | *IDH3G* |
| | *SLITRK4* | *IKBKG* |
| | *TGM4* | *JADE3* |
| | ***WNT7A*** | *NAA10* |
| | *ZIM2* | *NCAM2* |
| | *ZNF558* | *OCA2* |
| | | ***PCDHGA3*** |
| | | *PCDHGA5* |
| | | ***PLXNA3*** |
| | | *POTEF* |
| | | *RPL10* |
| | | *SLC10A3* |
| | | *SLC15A4* |
| | | *SLC16A2* |
| | | *SSR4* |
| | | *TMEM132D* |
| | | *TSPAN7* |
| | | *ZNF37A* |
| | | *ZNF41* |

**Table 5 Primary antibodies used for protein-based experiments.**

| Antibody | Species | Catalogue # | Dilution |
|---|---|---|---|
| MAP2 | mouse | Millipore mab3418a5 | 1:400 |
| VGLUT1 | Rabbit | Sigma 48-2400 | 1:1,000 |
| TUJ1 (Neuron-specific beta-III Tubulin) | Mouse | R&D Systems NL1195G | 1:200 |
| GFAP | Rabbit | Novus NB120-16997 | 1:200 |
| BrdU | Mouse | Invitrogen MA3-071 | 1:100 |
| H3.3 | Rabbit | Invitrogen PA5-29602 | 1:1,000 |
| H4 | Mouse | Cell Signaling #2935S | 1:500 |
| β-Actin | Mouse | Santa Cruz sc-69879 | 1:1,000 |
| SOX2 | Rabbit | Cell Signaling 35795 | 1:400 |
| Ki67 | Rabbit | Cell Signaling 9129 | 1:400 |
| PAX6 | Mouse | Sigma AMAb91372 | 1:100 |
| GABA | Rabbit | Sigma A2052 | 1:500 |

**Table 6 qPCR primers used to validate cell identity and validate RNA-seq results.**

| Gene | Forward primer | Reverse primer |
|---|---|---|
| *GAPDH* | GGCATCCTGGGCTACACTGA | GAGTGGGTGTCGCTGTTGAA |
| *H3-3B* | TCGTTGGGCGGTGCTGGTTTTT | TTAGGCCACTTCTCCGACCGCC |
| *GABRG3* | ATGCTACGCCAGCAAGAACAGC | GGCGAAGACAAACAGGAAGCAC |
| *DPP6* | GACCGACAGATGCCTAAAGTGG | TGTCGGTGAAGGTTGCTGGCTT |
| *GFAP* | CTGGAGAGGAAGATTGAGTCGC | ACGTCAAGCTCCACATGGACCT |
| *OCT4* | CCTGAAGCAGAAGAGGATCACC | AAAGCGGCAGATGGTCGTTTGG |
| *NANOG* | CTCCAACATCCTGAACCTCAGC | CGTCACACCATTGCTATTCTTCG |
| *SOX1* | GAGTGGAAGGTCATGTCCGAGG | CCTTCTTGAGCAGCGTCTTGGT |
| *NES* | TCAAGATGTCCCTCAGCCTGGA | AAGCTGAGGGAAGTCTTGGAGC |
| *MAP2* | AGGCTGTAGCAGTCCTGAAAGG | CTTCCTCCACTGTGACAGTCTG |
